# Mitochondrial H_2_O_2_ release does not directly cause genomic DNA damage

**DOI:** 10.1101/2023.03.29.534749

**Authors:** Daan M.K. van Soest, Paulien E. Polderman, Wytze T. F. den Toom, Susan Zwakenberg, Sasha de Henau, Boudewijn M. T. Burgering, Tobias B. Dansen

## Abstract

Reactive Oxygen Species (ROS) derived from mitochondrial respiration are frequently cited as a major source of genomic DNA damage and subsequent mutations that contribute to cancer development and aging. However, experimental evidence showing that ROS released by mitochondrial can directly damage nuclear DNA under (patho)physiological conditions has been largely lacking. In this study we modeled the effects of mitochondrial H_2_O_2_ release and compared this to H_2_O_2_ production at the nucleosomes in an untransformed human cell line. We used a chemogenetic approach to produce localized H_2_O_2_ and combined it with a new method we developed to directly quantify the amount of H_2_O_2_ produced. This enabled us to precisely investigate to what extent DNA damage occurs downstream of near- and supraphysiological amounts of localized H_2_O_2_ generation. Nuclear H_2_O_2_ production gives rise to DNA strand breaks, subsequent activation of the DNA damage response, cell cycle arrest and eventually senescence. Release of H_2_O_2_ from mitochondria on the other hand shows none of these effects, even at levels that are orders of magnitude higher than what mitochondria normally produce. Artificially high levels of mitochondrial H_2_O_2_ release do result in DNA strand breaks, but in parallel invariably cause ferroptosis-mediated cell death, preventing propagation of DNA damage-induced mutations. This study shows that H_2_O_2_ released from mitochondria is unlikely to directly damage genomic DNA, limiting its contribution to oncogenic transformation and aging.

## Introduction

Genomic DNA damage and subsequent mutation and selection is not only a driver of evolution but also of oncogenesis, as it can result in the acquisition of the hallmarks of cancer^1^. A tight control of genome integrity is therefore required to suppress malignant cell growth. The cellular DNA damage response (DDR), for instance, triggers a temporary cell cycle arrest to repair damaged DNA. Persistent DNA damage induces apopotosis or a permanent withdrawal from the cell cycle (senescence), which is thought to be both a mechanism for tumor suppression as well as a major cause of organismal aging^2^. DNA damage results from chemical modification, including DNA-base oxidation, which subsequently leads to mutagenesis because of mispairing during replication or inadequate repair. Enzymes that repair or prevent oxidative DNA lesions are found in all three domains of life (eukaryotes, bacteria & archaea), indicating that the prevention of genomic instability due to DNA oxidation is something many, if not all, organisms have to cope with^3-5^.

Mutagens that threaten DNA integrity do not only come from exogenous sources (like UV-light) but are also generated endogenously through cellular metabolism. Reactive Oxygen Species (ROS) are an example of potentially harmful metabolic products. ROS are generated at various intracellular sites in the form of superoxide (O_2_ ^•−^) or hydrogen peroxide (H_2_ O_2_)^6^. Importantly, both O_2_^•−^ and H_2_O_2_ are not reactive with DNA^7^. However, H_2_O_2_ can generate highly reactive hydroxyl radicals (^•^OH), catalyzed by ferrous iron (Fe^2+^) in the Fenton reaction, and the latter has been shown to be able to cause various oxidative DNA lesions *in vitro*^8,9^. The respiratory chain that drives mitochondrial ATP production is one of the major sites of metabolic ROS generation within the cell^10^. Respiratory chain derived ROS are generated in the form of O_2_^•−^, which are instable and very short-lived. O_2_^•−^ is rapidly converted to the more stable H_2_O_2_ by superoxide dismutases located in the cytoplasm and the mitochondrial inter-membrane space (SOD1) and the mitochondrial matrix (SOD2). The combined findings that exposure of cells to extracellular ROS triggers the DDR, that H_2_O_2_ (via ^•^OH) can oxidize DNA *in vitro*, and that mitochondria are one of the major intracellular suppliers of H_2_O_2_ is often extrapolated to the notion that ROS production and release by mitochondria leads to oxidative DNA damage and mutation^11-13^. This idea is also firmly rooted in the general public and is probably one of the foundations under the popular use of dietary antioxidant supplements. But evidence for direct oxidation of genomic (nuclear) DNA by mitochondria-derived ROS in live cells is largely lacking. As the diffusion range of ^•^OH is very limited due to its extreme reactivity, it would need to be formed close to the DNA to cause oxidative DNA-base modifications. This strongly suggests that DNA-base oxidation by mitochondrial ROS could only occur in case mitochondria-derived H_2_O_2_ would diffuse into the nucleus and form ^•^OH in proximity to the DNA. However, recent studies show that steep H_2_O_2_ gradients exist around the mitochondria due to efficient scavenging by peroxidases^14,15^, making it unlikely that H_2_O_2_ can act over distances large enough to reach the nucleus when released from mitochondria. Indeed, Cockayne syndrome-B (CS-B) mutated fibroblasts display elevated levels of mitochondrial ROS but this does not translate into increased nuclear DNA damage^16^. On the other hand, exogenously added H_2_O_2_ can trigger the DNA damage response^17^, but it is not clear how this compares to physiological continuous production of O_2_^•−^ and subsequently H_2_O_2_ by mitochondria in terms of achieved intracellular steady state and peak concentrations. Taken together, it remains unclear to what extent respiration-derived ROS contributes directly to DNA damage.

Recently, ectopic expression of the enzyme D-amino acid oxidase (DAAO) from the yeast *Rhodotorula gracilis* has been used to achieve titratable and sustained intracellular H_2_O_2_ production in live human cells as well as in *in vivo* rodent models^14,18,19^. H_2_O_2_ production is induced only upon administration of D-amino acids like D-Alanine, which are (largely, if not completely) absent from cultured cells. DAAO can be targeted to the organelle of choice by fusion to localization tags. H_2_O_2_ production by DAAO is often assessed using genetically encoded ratiometric fluorescent redox sensors like HyPer7^14,15^. Although these sensors are highly sensitive to H_2_O_2_, they report on the combined rate of oxidation and reduction, preventing direct quantification of absolute levels of H_2_O_2_ produced. To overcome this, we have setup a method to directly quantify DAAO dependent H_2_O_2_ production based on oxygen consumption rates, which allows for the careful titration of D-Ala levels that yield near and supraphysiological levels of H_2_O_2_ (van Soest et al., manuscript in preparation).

We set out to systematically analyze the localization-dependent cellular response to H_2_O_2_, making use of the chemogenetic production of H_2_O_2_ by ectopic expression DAAO. To mimic mitochondrial H_2_O_2_ release we fused DAAO to the cytoplasm-facing end of the targeting sequence of the outer mitochondrial membrane protein TOMM20 and stably expressed it in RPE1-hTert cells. RPE1 cells expressing nucleosome-targeted DAAO serve as a positive control for what happens when H_2_O_2_ reaches the nuclear DNA. Careful titration of the substrate for DAAO enables the continuous production of near and supra-physiological H_2_O_2_ levels. Using this approach, we show that H_2_O_2_ released from mitochondria does not induce direct damage to the nuclear DNA when produced at levels likely achievable by respiration. In contrast, H_2_O_2_ generation in close proximity to the nuclear DNA causes DNA strand breaks and subsequent activation of the DNA damage response (DDR), resulting in cell cycle arrest and senescence, similar to as what has been described for cells exposed to high levels of exogenous ROS. Based on these observations we conclude that mitochondrial respiration-derived ROS is probably not a major factor in the induction of genomic mutations, at least in non-transformed cells.

## Results

### Stable expression of D-amino acid oxidase (DAAO) in RPE1-hTert enables localized intracellular H_2_O_2_ production

The various effects of ROS on cells have mostly been studied by applying a bolus (typically 10-200 µM) of H_2_O_2_ to the tissue culture media. Although such treatments certainly induce oxidative stress, it does not recapitulate the continuous ROS production (i.e. O_2_^•−^ and subsequently H_2_O_2_) by mitochondria, which occurs at far lower levels. We therefore made use of chemogenetic production of H_2_O_2_ using localized DAAO expression. For this study we generated monoclonal RPE1^hTERT^ cell lines stably expressing mScarlet-I-DAAO localized to two different sites (fig. 1A). H2B-mScarlet-I-DAAO localizes to nucleosomes to study the effects of H_2_O_2_ when it is produced in close proximity to DNA. TOM20-mScarlet-I-DAAO localizes to the cytoplasmic side of the outer mitochondrial membrane and hence mimics the release of H_2_O_2_ by mitochondria. RPE1-hTert cells were chosen as this human cell line is untransformed and the signal transduction pathways relevant for cell cycle regulation and the DNA damage response are fully functional. We will refer to these cell lines from now on as RPE1-hTert-DAAO^H2B^ and RPE1-hTert-DAAO^TOM20^. Imaging of mScarlet-I confirms the correct localization of DAAO by colocalization with DNA (Hoechst 33342 staining) for RPE1-HTERT-DAAO^H2B^ cells and mitochondria (MitoTracker Green staining) for RPE1-hTert-DAAO^TOM20^ cells respectively (fig. 1B).

**Figure 1:**
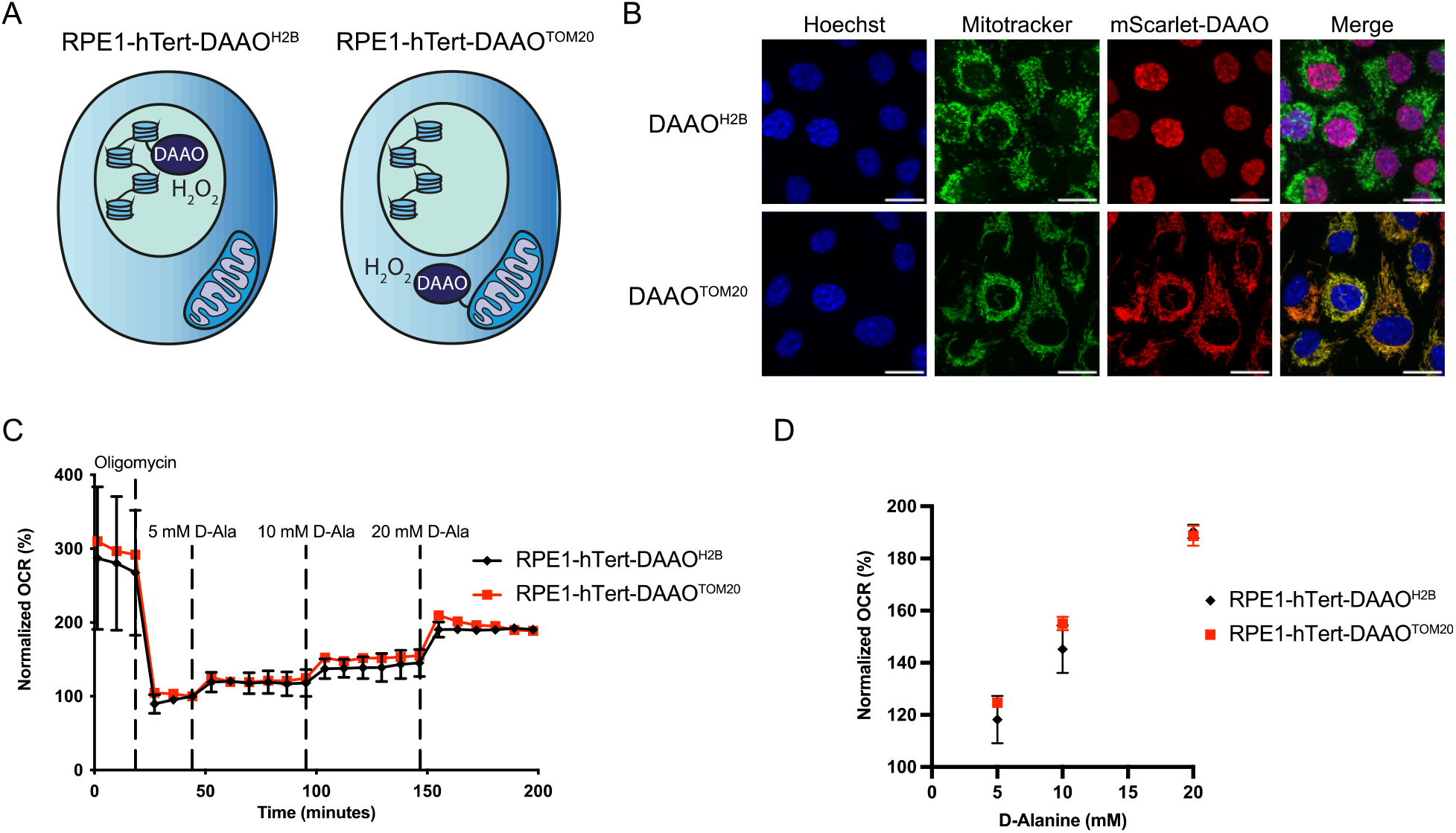
Localized expression of DAAO enables intracellular H_2_O_2_ production. (A) Schematic representation of the generated RPE1^hTERT^ cells expressing H2B-mScarlet-I-DAAO or TOM20-mScarlet-I-DAAO to induce localized intracellular H_2_O_2_ production. (B) Immunofluorescence confocal images of cell lines depicted in (A) showing colocalization of DAAO^H2B^ and DNA (Hoechst), and colocalization of DAAO^TOM20^ with mitochondria (mitotracker green). Scale bar equals 10 μm. (C) Oxygen consumption rate (OCR) upon D-Ala administration as a readout for enzymatic activity of ectopically expressed DAAO. The increase in OCR show that both lines produce roughly similar amounts of H_2_O_2_ upon addition of D-Ala. Data is normalized to OCR at the last timepoint before D-Ala injection. Error bars represent SD of 5 technical replicates. (D) Normalized OCR from (C) plotted against D-ala concentration, showing that DAAO activity increases proportionally with the D-Ala concentration used.

Total DAAO activity in these cell lines is likely determined by a combination of the level of DAAO expression, folding, accessibility, cofactor (FAD) availability as well as substrate (D-amino acid) levels, uptake, and diffusion^20^. Because DAAO uses equimolar amounts of molecular oxygen to produce H_2_O_2_, we assessed DAAO activity by measuring the oxygen consumption rate (OCR) in response to titration with [D-Ala]. Mitochondrial respiration of RPE1-HTERT-DAAO^H2B^ and RPE1-hTert-DAAO^TOM20^ cells was inhibited by oligomycin prior to D-Ala addition in order to minimize differences in oxygen consumption that stem from metabolic respiration and to be able to compare DAAO-derived H_2_O_2_ to mitochondrial OCR. In this way monoclonal RPE1-hTert-DAAO^TOM20^ and RPE1-HTERT-DAAO^H2B^ lines with similar DAAO activities were identified and selected (Figure 1 C,D). Thus, observed differences between these lines upon DAAO activation are the result of differential localization of H_2_O_2_ production and not of total oxidative burden. Both cell lines consume an amount of oxygen equivalent to roughly 10% of basal mitochondrial respiration at 5 mM D-alanine. It is often stated that about 2% of oxygen escapes the mitochondrial ETC as O_2_^•−^, although this number is based on maximum O_2_^•−^ production by complex III under saturating oxygen and substrate conditions using isolated mitochondria, whereas O_2_^•−^ production by the ETC in live cells has been suggested to be ten times lower^21^. Furthermore, it takes two molecules of O_2_^•−^ to produce one molecule of H_2_O_2_, and not all H_2_O_2_ will be released from the matrix and the IMS into the cytosol due to scavenging by for instance PRDX3. Hence, at 5 mM of D-Ala, mitochondrial H_2_O_2_ release as modeled by the RPE1-hTert-DAAO^TOM20^ cell line is already far above what can be expected to occur under (patho)physiological conditions.

To investigate the levels of DAAO activity (via a different approach), we performed non-reducing western blots for the H_2_O_2_ scavenger peroxiredoxin 2 (PRDX2) (Sup. fig.1). PRDX2 is found in the cytosol as well as the nucleus of cells. Upon oxidation of its H_2_O_2_-sensitive cysteines PRDX2 forms homodimers, which can be detected by an upward band shift on a non-reducing gel. Indeed, increasing amounts of D-Alanine resulted in increased dimerization of PRDX2. The extent of dimerization was similar between H2B-DAAO and TOM20-DAAO cells, which is in line with OCR measurements (fig. 1C,D). At high levels of H_2_O_2_, peroxiredoxins can become inactivated through overoxidation (PRDXSO_2/3_). In contrast to exogenous H_2_O_2_ treatment, no overoxidation of Prdx was detected upon DAAO activation at viable levels. This indicates that continues intracellular H_2_O_2_ production evokes a very different cellular redox response as compared to a bolus of exogenous H_2_O_2_.

### Diffusion of locally produced H_2_O_2_ is limited

For respiration-derived ROS to directly inflict damage to genomic DNA it would need to diffuse out of mitochondria, across the cytoplasm and over the nuclear membrane. O_2_^•−^ has a very limited diffusion range due to its low membrane permeability^22^ and rapid conversion to H_2_O_2_ by Superoxide dismutases^23^. H_2_O_2_ has been suggested to be stable enough to travel some distance in biological systems, although the exact range is debated. We assessed the diffusion range of H_2_O_2_ produced by DAAO at the outer mitochondrial membrane and at the nucleosome using live fluorescence microscopy of the ultrasensitive, ratiometric H_2_O_2_ sensor HyPer7 stably expressed in the nucleus (NLS-HyPer7) and the cytoplasm (NES-HyPer7). Oxidation of cytosolic HyPer7 was readily detected at low [D-Ala] in RPE1-hTert-DAAO^TOM20^ cells, and nuclear HyPer7 rapidly responded to low [D-Ala] in RPE^H2B-DAAO^ cells, showing that HyPer7 is oxidized when in close proximity to the site of H_2_O_2_ production (fig. 2A,C). In contrast, low/medium levels of H_2_O_2_ produced by mitochondrial membrane-localized DAAO did not result in oxidation of nuclear HyPer7 (fig. 2B). Only at very high DAAO substrate levels (> 10 mM D-Ala) an increase in NLS-HyPer7 was detected in RPE1-hTert-DAAO^TOM20^ cells. However, cells were not able to survive this amount of H_2_O_2_ production for prolonged duration (fig. 2E & sup. fig. 2). A similar trend was observed for the combination of NES-HyPer7 in RPE1-HTERT-DAAO^H2B^ cells: H_2_O_2_ is only detected outside the nucleus when high [D-Ala] are supplied (also > 10 mM) (fig. 2C-D). Similarly, these high [D-Ala] lead to levels of H_2_O_2_ production that are not compatible with cell survival (fig. 2E). These observations are in line with earlier studies that suggested that in cells H_2_O_2_ is mostly confined to the site of production. Furthermore, when H_2_O_2_ is produced at levels high enough to diffuse from the cytoplasm into the nucleus (or vice versa), this is incompatible with cell survival. Note that based on our OCR experiments (fig. 1C-D) it can be estimated that the amount of H_2_O_2_ production needed to achieve diffusion from the outer mitochondrial membrane into the nucleus corresponds to ∼25% of total oxygen used in mitochondrial respiration, which is difficult to imagine occurring under (patho)physiological conditions. Based on these observations and considerations we propose that it is highly unlikely that under physiological circumstances respiration-derived ROS can diffuse into the nucleus and directly induce oxidative damage to genomic DNA that could lead to mutations.

**Figure 2:**
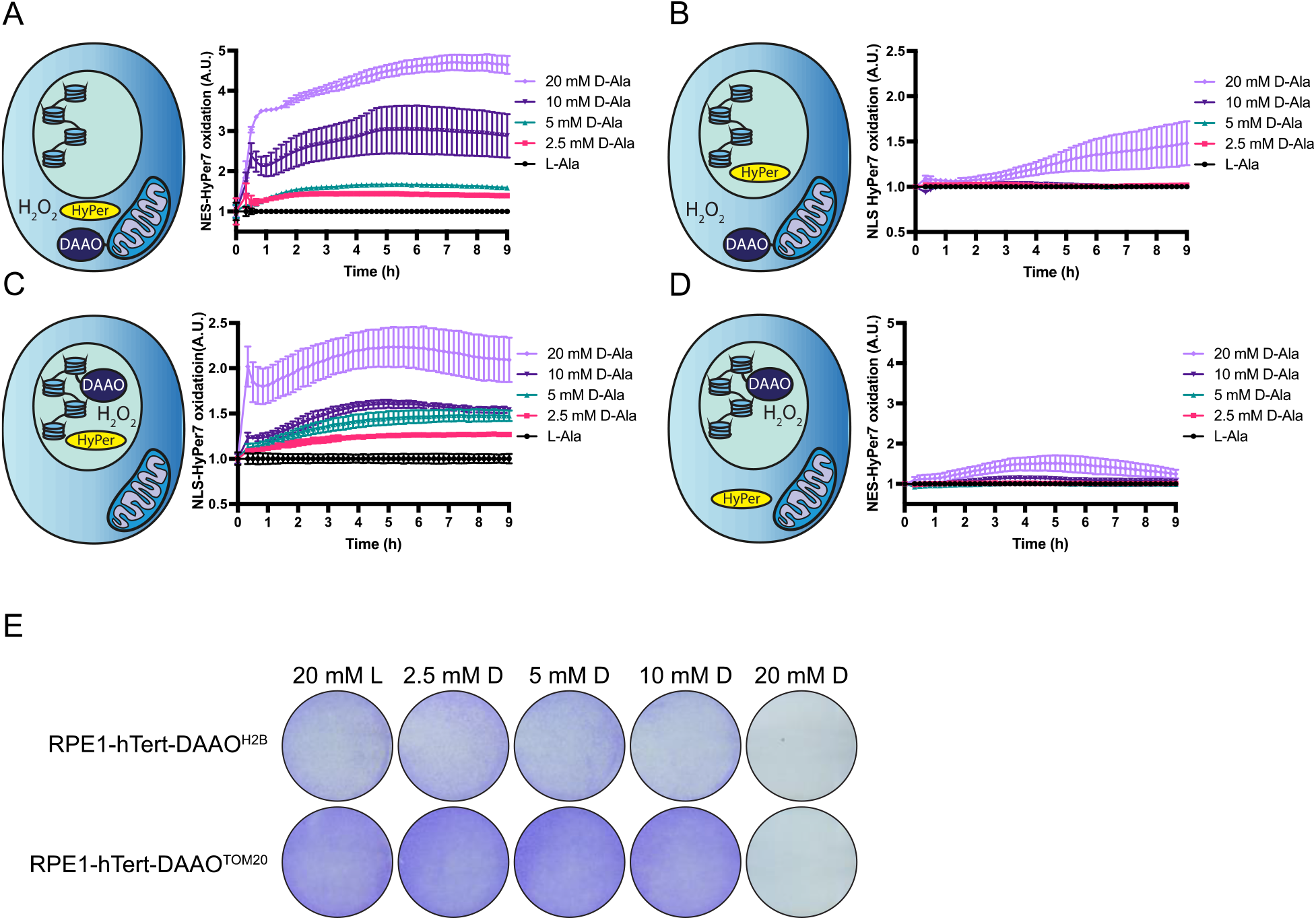
H_2_O_2_ released by mitochondria is not detected in the nucleus at levels that are compatible with cell survival. (A-D) Intracellular measurements using the ratiometric H_2_O_2_ sensor HyPer7 to determine diffusion of DAAO-produced H_2_O_2_. Cytosolically localized NES-HyPer7 is already oxidized at the lowest concentration of D-Ala used (2.5 mM) in RPE1-hTert-DAAO^Tom20^ cells. Nuclear located NLS-HyPer7 oxidation is only detected above 10mM D-Ala in this cell line (B). In contrast, NLS-HyPer7 oxidation is already detected at 2.5 mM D-Ala in RPE1-hTert-DAAO^H2B^ cells (C), while NES-HyPer7 oxidation is only detected above 10 mM D-Ala. (E) Crystal violet staining of DAAO lines after 24h L/D-Ala treatment. Both RPE1-hTert-DAAO^H2B^ and RPE1-hTert-DAAO^Tom20^ cells do not survive D-Ala treatments above 10 mM D-Ala, suggesting cell survival is only compatible with levels of DAAO activation where H_2_O_2_ remains close to its site of production.

### Mitochondrial H_2_O_2_ release does not induce genomic DNA damage

H_2_O_2_ derived from mitochondria may not travel to the nucleus and cause DNA damage by direct oxidation, but in principle DNA damage downstream of H_2_O_2_ could occur through mechanisms that would not require diffusion of H_2_O_2_ into the nucleus. Firstly, cytoplasmic H_2_O_2_ could lead to oxidation of the nucleotide pool and subsequent incorporation of these oxidized nucleotides in the DNA during replication. Secondly, cytoplasmic H_2_O_2_ may hamper the fidelity or activity of proteins involved in DNA repair and replication by direct or indirect oxidation. The DNA repair protein RAD51 for instance has been shown to be inhibited upon Cysteine oxidation. Treatment of cells with a high dose of exogenous H_2_O_2_ has been shown to activate both ataxia telangiectasia and RAD3-related protein (ATR) and ataxia telangiectasia mutated protein (ATM) dependent DNA damage response (DDR), indicative of single and double strand DNA breaks respectively. Indeed, the repair of 8-oxo-dG in genomic DNA proceeds through a transient single strand and may even lead to double strand breaks^24^. Therefore, we monitored whether the DDR was activated in RPE1-hTert-DAAO^TOM20^ and RPE1-HTERT-DAAO^H2B^ lines upon D-Ala treatment in. Indeed, induction of nuclear H_2_O_2_ production resulted in a clear activation of ATR and ATM, indicated by phosphorylation of their downstream targets checkpoint kinases CHK1 and CHK2 respectively (fig. 3A-B). This coincided with the stabilization of p53 and subsequent transcription of the cell cycle inhibitor p21. Knockout of p53 resulted in even higher levels of pCHK1, pCHK2 and γH2AX, likely due to a sustained DDR because of impaired DNA repair. However, without p53 this was not translated into transcription of p21. In contrast to H_2_O_2_ produced at the nucleosome, mimicking the release of mitochondrial H_2_O_2_ by TOM20-DAAO did not result in activation of the DDR. Even in the absence of p53, no increase in CHK1 or CHK2 phosphorylation or γH2AX levels was observed.

**Figure 3:**
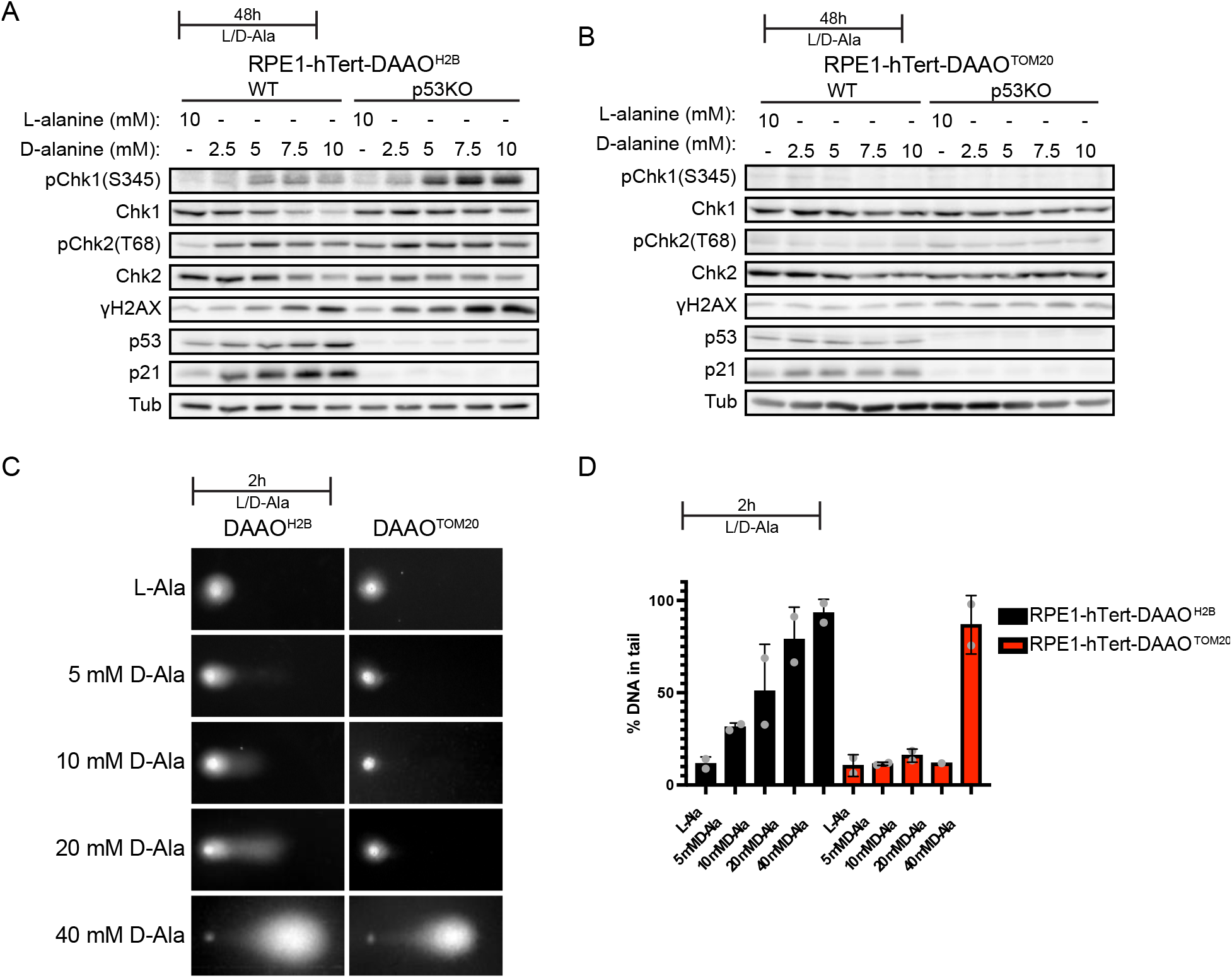
DNA strand breaks and subsequent activation of the DNA damage response can be caused by nuclear H_2_O_2_ but not by mitochondrial H_2_O_2_ release. (A-B) Western Blot for components of the DNA damage response in WT or p53KO RPE1-hTert-DAAO^H2B^ and RPE1-hTert-DAAO^TOM20^ cells treated for 48h with L-or D-alanine. Nuclear H_2_O_2_ production results in activation of the DNA damage response (pchk1, pchk2, γH2AX) and subsequent p53 stabilization and p21 expression. Activation of DNA damage response is even stronger in the absence of p53. In contrast, H_2_O_2_ produced at mitochondria by DAAO^TOM20^ does not activate the DNA damage response. (C) Representative images of comet assay with RPE1-hTert-DAAO^H2B^ and RPE1-hTert-DAAO^TOM20^ cells treated for 2h with L-or D-Ala. H_2_O_2_ production at mitochondria do not result in DNA strand breaks at levels that are compatible with survival (≤10 mM D-Ala). However, H_2_O_2_ production close to the DNA does give rise to DNA strand breaks, as shown by the increase of DNA in the comet tail. (D) Quantification of percentage of DNA in comet tail of comets shown in A. The average from 2 biological repeats is shown, each biological repeat containing 50 analyzed comets per condition. Error bars depict SD.

Activation of the DDR is an indirect measurement of DNA damage. To test whether DNA single and double strand breaks occur downstream of localized H_2_O_2_ production in a more direct way, we performed alkaline comet assays. RPE1-HTERT-DAAO^H2B^ lines showed a dose-dependent increase in DNA breaks upon D-Ala treatment, indicating that H_2_O_2_ indeed is capable of inducing DNA strand breaks (fig. 3 C-D). Note that DNA breaks can be detected at levels of DAAO-induced H_2_O_2_ that are not overtly toxic (e.g., at <10 mM D-Ala), suggesting that this amount of damage is still repairable. In contrast, no increase in DNA strand breaks was detected when RPE1-hTert-DAAO^TOM20^ were treated with [D-Ala] compatible with cell survival (fig. 2E). At 40 mM D-Ala DNA strand breaks can be detected in RPE1-hTert-DAAO^TOM20^ cells, but it is unlikely that these levels of H_2_O_2_ can be achieved physiologically, as this likely corresponds to an amount of H_2_O_2_ production two orders of magnitude higher than what mitochondria normally produce (fig. 1C). Besides, this level of H_2_O_2_ production at the mitochondria invariably kills the cells, which precludes mutation accumulation down the lineage (fig. 2E & sup. fig 2).

These experiments show that H_2_O_2_ produced in close proximity to the DNA can indeed trigger DNA breaks and a subsequent DNA damage response. H_2_O_2_ released by mitochondria is not able to induce these phenotypes at levels compatible with cellular survival, suggesting that mitochondria-derived H_2_O_2_ also does not indirectly contribute to oxidative DNA damage.

### A p53-dependent cell cycle arrest and senescence is induced by H_2_O_2_ production in proximity to DNA but not by mitochondrial H_2_O_2_ release

Exogenous addition of H_2_O_2_ induces a cell cycle arrest in RPE1-hTert cells^25^. To investigate whether localized H_2_O_2_ production can also trigger a cell cycle arrest, and whether that depends on its site of production, we performed a BrdU incorporation assay. RPE1-hTert-DAAO^TOM20^ and RPE1-HTERT-DAAO^H2B^ cells were treated for 48h with L-or D-Ala, followed by incubation in media containing BrdU (and no L-or D-Ala) for 24h to label cycling cells. BrdU positive and negative cells were assessed by flow cytometry. In line with our observation that H_2_O_2_ production at the nucleosome triggers p53 stabilization and activity, H_2_O_2_ production in RPE1-HTERT-DAAO^H2B^ cells resulted in a clear loss of proliferative capacity upon treatment with D-Ala (fig. 4A). H_2_O_2_ release from mitochondria on the other hand failed to induce a cell cycle arrest at any concentration that did not result in cell death. In line with the observed stabilization and activation of p53 upon H_2_O_2_ production at the nucleosome (fig. 3A), the observed arrest in RPE1-HTERT-DAAO^H2B^ cells was indeed p53 dependent (fig. 4B). We then assessed the ploidy of the arrested cells by cell cycle profiling by flow cytometry. RPE1-HTERT-DAAO^H2B^ cells with 4N DNA accumulated upon induction of H_2_O_2_ production, whereas the cell cycle profile of RPE1-hTert-DAAO^TOM20^ cells remained unchanged (fig. 4C&D). The induction of the arrest with a ploidy of 4N in RPE1-HTERT-DAAO^H2B^ cells was also largely p53 dependent (fig. 4F). To discriminate cells arrested with 2N DNA from the majority of cycling cells that reside transiently in G1 we performed a similar experiment but trapped cycling cells in mitosis using the spindle poison nocodazole during the last 16 hours of D-Ala treatment. There was no sign of a cell cycle arrest with 2N DNA in either cell line (fig. 4E).

**Figure 4:**
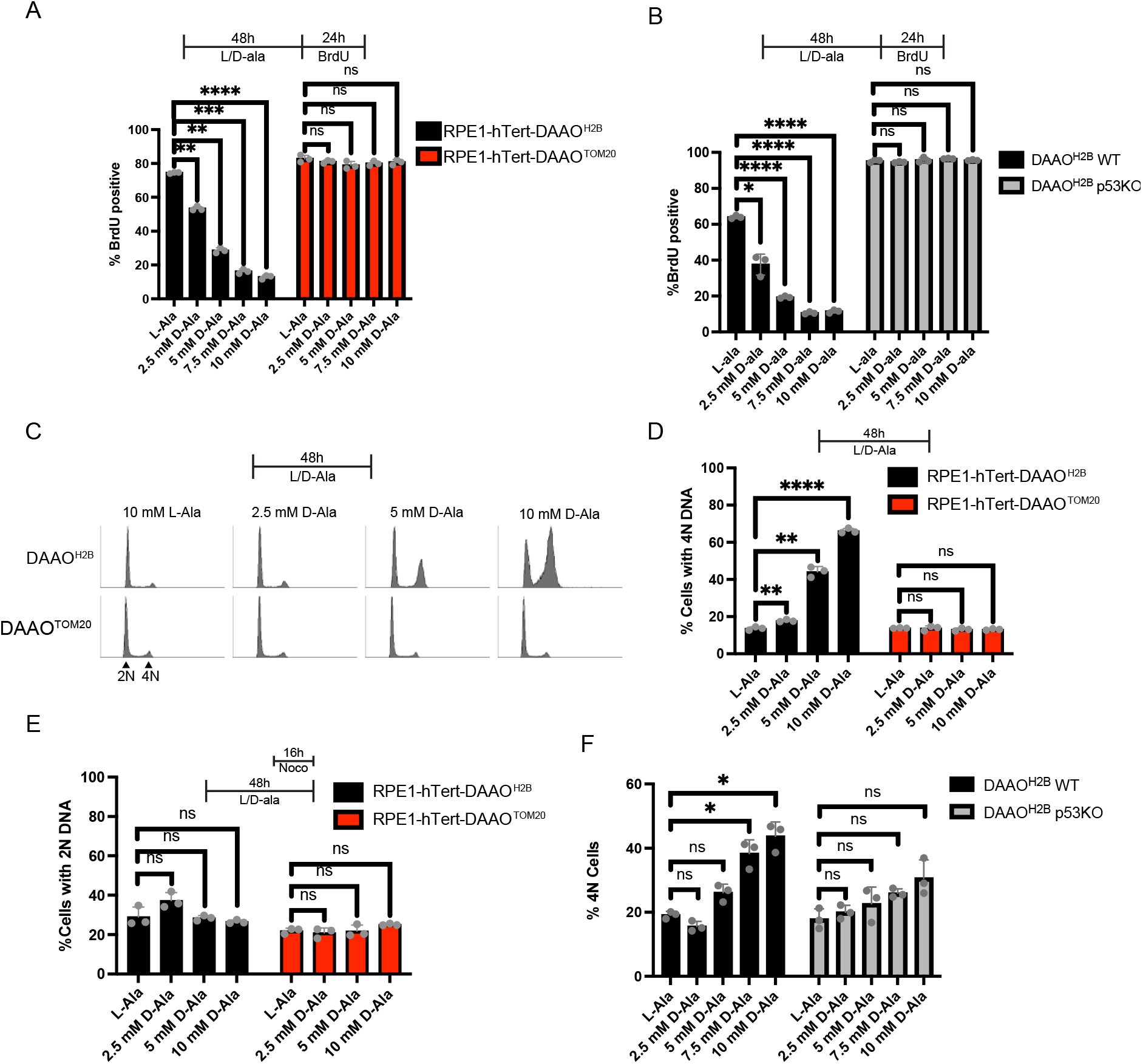
Nuclear H_2_O_2_ causes a p53-dependent 4N cell cycle arrest, while mitochondrial H_2_O_2_ release does not. (A) Quantification of BrdU assay. Cells were treated for 48h with L-or D-Ala and passaged to larger dishes to prevent contact inhibition. Cells were subsequently cultured for 24h in media containing BrdU. Cells were then fixed, stained for BrdU and analysed by Flow cytometry. Induction of nuclear H_2_O_2_ clearly decreases the amount of cycling cells, while mimicking mitochondrial H_2_O_2_ release by DAAO^TOM20^ activation does not. Dots represent 3 biological replicates; Each replicate encompasses analysis of roughly 10.000 cells. Statistical test performed is Welch ANOVA test + Dunnett’s T3 multiple comparison test (**p≤0.005, *** p≤0.0005, ****p≤0.0001). D-Ala treatments in RPE1-hTert-DAAO^TOM20^ cells did not yield statistically different results from the L-Ala control. (B) Quantification of BrdU assay. RPE1-hTert-DAAO^H2B^ WT or p53KO cells were treated for 48h with L-or D-Ala and passaged to larger dishes to prevent contact inhibition. Cells were subsequently cultured for 24h in medium containing BrdU. Cells were then fixed, stained for BrdU and analysed by Flow cytometry. Cell cycle arrest induced by DAAO^H2B^ is dependent on p53, as p53KO cells do not increase in BrdU negative cells. Dots represent 3 biological replicates; Each replicate encompasses analysis of roughly 10.000 cells. Statistical test performed is Welch ANOVA test + Dunnett’s T3 multiple comparison test (*p≤0.05, ****p≤0.0001). (C) DNA profile by PI staining and subsequent FACS analysis of RPE1-hTert-DAAO^H2B^ and RPE1-hTert-DAAO^TOM20^ cells. Cells were treated for 48h with L-or D-Ala. The induction of nuclear H_2_O_2_ production by D-Ala treatment of RPE1-hTert-DAAO^H2B^ cells causes a clear 4N cell cycle arrest, while this is absent when mimicking mitochondrial H_2_O_2_ release in RPE1-hTert-DAAO^TOM20^ cells. (D) Quantification of DNA profile in (A), showing the percentage of cells with a 4N DNA content. Dots represent 3 biological replicates; Each replicate encompasses analysis of roughly 10.000 cells. Statistical test performed is Welch ANOVA test + Dunnett’s T3 multiple comparison test (**p≤0.005, ****p≤0.0001). All D-Ala treatments in RPE1-hTert-DAAO^TOM20^ cells were not statistically different from the L-Ala control. (E) Quantification of DNA profiles, showing the percentage of cells with a 2N DNA content. RPE1-hTert-DAAO^H2B^ and RPE1-hTert-DAAO^TOM20^ cells were treated for 48h with L-or D-Alanine. The last 16h of treatment, nocodazole was added to block cells in metaphase to help visualize a potential 2N cell cycle arrest. No increase in the percentage of 2N cells is observed upon D-Ala addition in both lines, demonstrating there was no 2N cell cycle arrest initiated. Dots represent 3 biological replicates; Each replicate encompasses analysis of roughly 10.000 cells. (F) Quantification of DNA profiles by PI staining and subsequent FACS analysis. RPE1-hTert-DAAO^H2B^ WT and p53KO cells were treated for 48h with L-or D-Ala. Graph shows the percentage of cells with a 4N DNA content. Dots represent 3 biological replicates; Each replicate encompasses analysis of roughly 10.000 cells. Statistical test performed is Welch ANOVA test + Dunnett’s T3 multiple comparison test (*p≤0.05).

An arrest with 4N DNA does not necessarily mean that the cells are arrested in G2 or M phase of the cell cycle: previous studies have shown that hydrogen peroxide treatment can induce mitotic bypass after which cells permanently exit the cell cycle with 4N DNA^26-28^ The observed arrest with 4N DNA in RPE1-HTERT-DAAO^H2B^ cells was maintained despite D-Ala being washed away before BrdU incorporation and we tested whether these cells indeed became senescent. To investigate this, cells were kept in culture for an additional 5 days after treatment to allow expression of senescence markers. RPE1-HTERT-DAAO^H2B^ cells indeed gained senescence associated-β-galactosidase activity upon treatment with D-Ala (fig. 5A). Accordingly, immunofluorescence microscopy indicated that RPE1-HTERT-DAAO^H2B^ cells accumulated p21 and lost LaminB1, and displayed senescence-associated heterochromatin foci (SAHFs), all of which are makers of senescent cells^29^ (fig. 5B-C, S3). RPE1-hTert-DAAO^TOM20^ cells on the other hand showed no sign of senescence induction. In summary, increased H_2_O_2_ production near the DNA causes p53-dependent cell cycle arrest with 4N DNA and senescence, whereas H_2_O_2_ release from mitochondria does not.

**Figure 5:**
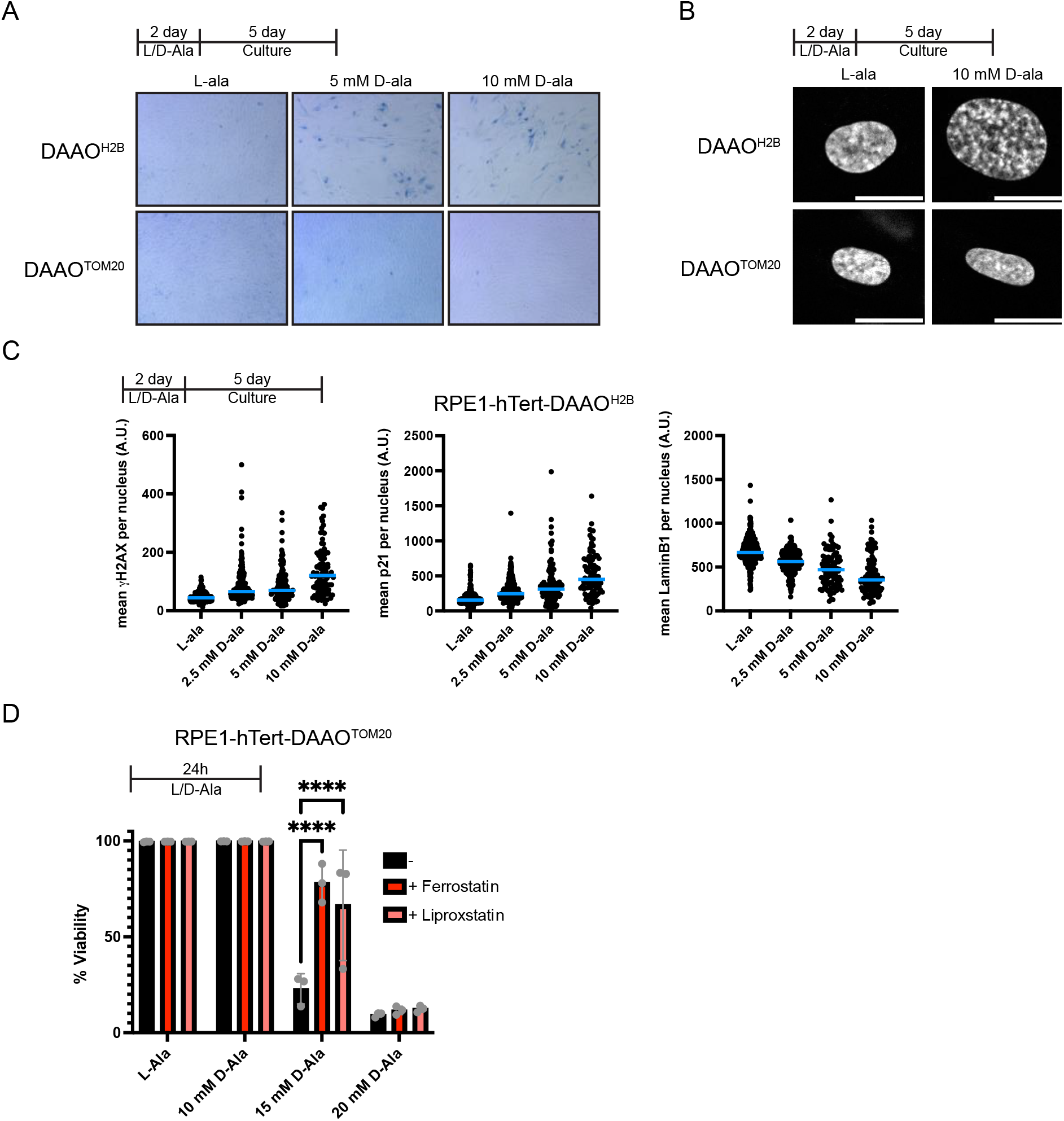
Nuclear H_2_O_2_ production causes senescence, while senescence induction by excessive H_2_O_2_ release is prevented by ferroptosis induction. (A) Senescence Associated-β-gal staining of RPE1-hTert-DAAO^H2B^ and RPE1-hTert-DAAO^TOM20^ cells. Cells were treated for 48h with L-or D-Ala, after which they were cultured for an additional 5 days to allow for expression of senescence markers. H2B-DAAO cells become positive for the staining upon D-Ala addition, while TOM20-DAAO cells remain negative. (B) DAPI staining of RPE1-hTert-DAAO^H2B^ and RPE1-hTert-DAAO^TOM20^ cells. Cells were treated for 48h with L-or D-Ala, after which they were cultured for an additional 5 days to allow for expression of senescence markers. RPE1-hTert-DAAO^H2B^ cells undergo chromatin restructuring in response to D-Ala treatment, demonstrating a phenotype reminiscent of senescence-associated heterochromatin foci (SAHFs). This remains absent in RPE1-hTert-DAAO^TOM20^ cells. (c) Quantification of IF stainings for p21, laminB1 and yH2AX. Average signal per nuclei is shown. Cells were treated for 48h with L-or D-Ala, after which they were cultured for an additional 5 days to allow expression of senescence markers. Induction of nuclear H_2_O_2_ results in a decrease in nuclear laminb1 signal and an increase in nuclear p21 intensity, both markers of senescence. RPE1-hTert-DAAO^H2B^ cells also show an increase in H2AX signal upon D-Ala addition, which indicates DNA damage occurred. (D) Quantification of cell viability by PI exclusion in RPE1-hTert-DAAO^TOM20^ cells. Dots represent 3 biological replicates. Per replicate, 10.000 cells were analyzed by FACS. Massive cell death occurs at 15 mM D-Ala, which can be rescued by ferroptosis inhibitors ferrostatin and liproxstatin. 2-way Anova + Dunett’s multiple comparison test ****=0.0001

### Supraphysiological mitochochondrial H_2_O_2_ release induces cell death by ferroptosis

H_2_O_2_ production at the mitochondrial outer membrane in RPE1-hTert-DAAO^TOM20^ cells does not trigger a cell cycle arrest (fig. 4A), but high concentrations of D-Ala these cells did no longer grow out (fig. 2E, S2). We wondered whether these cells succumb by ferroptosis, which can be triggered when H_2_O_2_ forms hydroxyl radicals in the presence of iron. Indeed, pre-treatment with the ferroptosis inhibitors ferrostatin or liproxstatin could partially rescue cell death, although at higher [D-Ala] they still died (Fig. 5D). In the current experimental setup, the induction of ferroptosis occurs at levels of H_2_O_2_ production at the mitochondrial outer membrane that are high enough to be picked up by the HyPer7 sensor in the nucleus (Fig 2), and hence outcompete the cytoplasmic antioxidant capacity. Collectively, our data suggest that mitochondria-derived H_2_O_2_ does not directly contribute genomic DNA damage because it does not reach the nucleus under normal conditions. Even if sufficient H_2_O_2_ could be produced by mitochondria that it could reach and damage nuclear DNA, which seems unlikely based on the comparison with mitochondrial OCR (Fig 1C), the cells die by ferroptosis, preventing mutation accumulation and propagation (fig 6).

**Figure 6:**
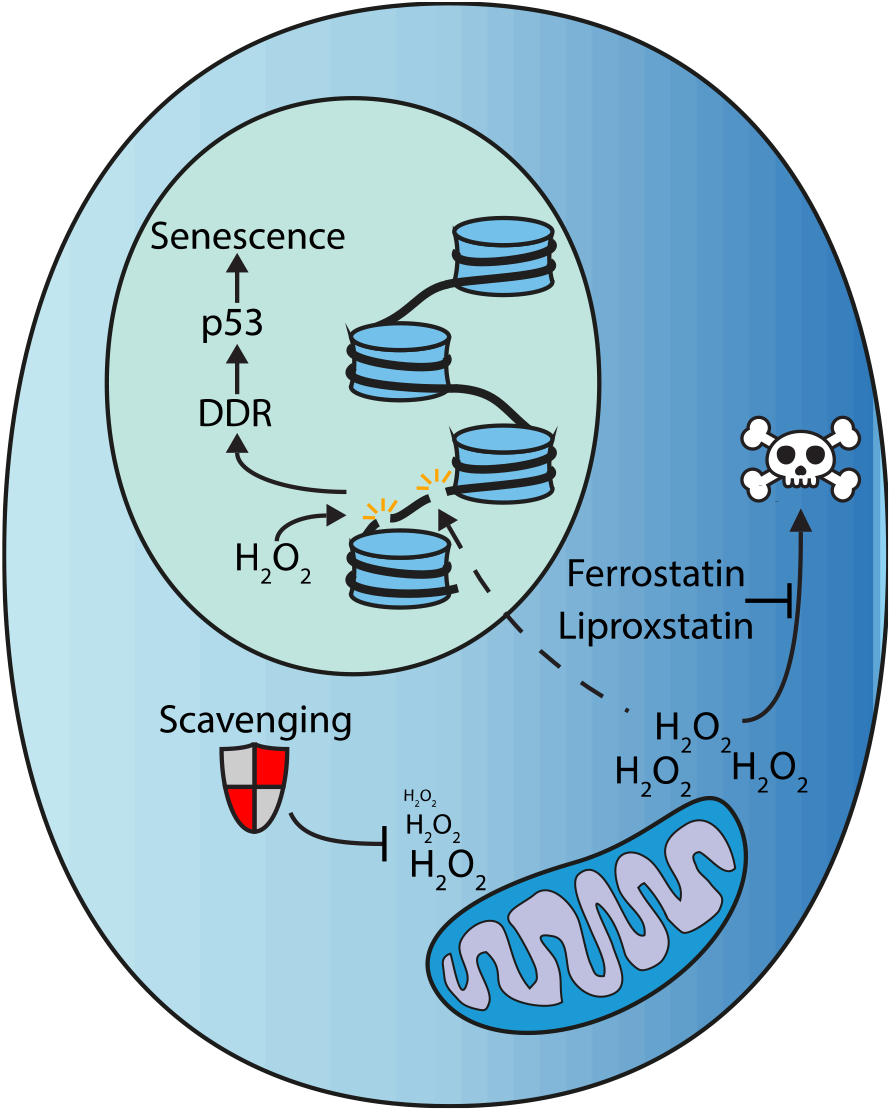
Mitochondrial H_2_O_2_ release does not directly damage nuclear DNA. Graphical abstract of our findings. Continuous elevated H_2_O_2_ levels in close vicinity to the DNA can cause DNA strand breaks, resulting in activation of the DNA damage response and ultimately senescence. In contrast, H_2_O_2_ released from mitochondria is not able to reach the nuclear DNA at physiological levels, and therefore does not directly contribute to nuclear DNA damage. Artificially high levels of mitochondrial H_2_O_2_ release does result in DNA strand breaks, but in parallel causes ferroptosis-mediated cell death, preventing eventual mutations to be propagated.

## Discussion

DNA can be oxidized by certain reactive oxygen species (mainly ^•^OH) *in vitro*^*8,9*^, and the DDR is triggered in cells exposed to supraphysiological amounts of exogenous H_2_O_2_^17^. Oxidized DNA bases, predominantly 8-oxo-dG, can be detected in patients with various diseases as well as in healthy subjects. 8-oxo-dG in the DNA can mispair with adenine, leading to C>A transversions after replication: a COSMIC Single Base Substitution (SBS) signature that can be found in several types of cancer^30^. In literature, these separate observations are often combined and extrapolated as if the (elevated) production and release of ROS by mitochondria can be directly translated to increased DNA oxidation and subsequent mutagenesis^11-13^. Based on a similar rationale, ROS derived from oxygen dependent mitochondrial ATP production is frequently implicated in cancer initiation, progression and treatment resistance, but also in aging in general. Unambiguous evidence for the *direct* oxidation and subsequent mutation of genomic DNA by ROS released from mitochondria at physiological levels however, is difficult to find.

Recent studies have employed sustained localized H_2_O_2_ production by DAAO to study the diffusion range of H_2_O_2_ and concluded that this is very limited due to highly efficient scavenging by abundant peroxidases^14,15,31^. But most of these studies used oncogenic transformed cell lines with a compromised p53 and DDR, which are therefore less suited to study the effect of mitochondrial H_2_O_2_ release on DNA damage and DNA damage-mediated cell cycle arrest. Here we used this chemogenetic approach to induce local intracellular H_2_O_2_ production at the mitochondria or at nucleosomes to investigate the potential role of intracellular H_2_O_2_ production in DNA damage in untransformed human RPE1-hTert cells. H_2_O_2_ generated at the nucleosomes indeed induced DNA strand breaks, resulting in activation of the DDR and subsequent p53-dependent cell cycle arrest and senescence. Mitochondrial H_2_O_2_ release did not result in a detectable increase of H_2_O_2_ in the nucleus, unless H_2_O_2_ was produced at levels that are one or two orders of magnitudes higher than what would likely be achievable physiologically. Furthermore, mitochondrial H_2_O_2_ release did not cause DNA strand breaks and activation of the DNA damage response at physiologically levels. Even at levels of mitochondrial H_2_O_2_ release that are equivalent to at least 10 to 100-fold the amount of ROS that is reportedly normally produced within mitochondria, cells maintain the capacity to proliferate without signs of DNA damage. Only when H_2_O_2_ production by DAAO^TOM20^ at the mitochondria is boosted even further, so that it can be detected in the nucleus by Hyper7 (fig. 2A), it results DNA strand breaks as shown in the comet assay (fig. 3C-D). However, and importantly, these levels of mitochondrial H_2_O_2_ production and release are unlikely to occur as a result of mitochondrial respiration and cells are not able to maintain redox homeostasis and die through ferroptosis.

We cannot completely exclude the possibility that release of mitochondrial H_2_O_2_ can indirectly result in DNA mutations. Free nucleotides, synthesized in the cytoplasm, could be oxidized by ^•^OH and subsequently incorporated in the DNA during S-phase. The presence of the enzyme MutT Homologue 1 (MTH1, also known as Nudix Hydrolase 1 (NUDT 1)), which sanitizes the oxidized free nucleotide pool is an indication that this could be a feasible scenario^32^. However, repair of 8-oxo-dG through BER proceeds through single stranded DNA intermediates, and we found no evidence of enhanced DNA strand breaks in the comet assay nor activation of ATM, ATR or p53 and cell cycle arrest in response to mitochondrial H_2_O_2_ release. We therefore conclude that, at physiologically levels, the release of ROS by mitochondria is not a significant source of direct DNA oxidation or indirect DNA damage and is therefore not likely to contribute considerably to cancer initiation and aging.

Although mitochondrial H_2_O_2_ *release*, as modeled here by TOM20-DAAO, may not contribute directly to DNA oxidation, H_2_O_2_-dependent and cysteine-thiol-based redox signaling is suggested to directly regulate DNA damage response and repair enzymes, as has been shown for ATM^33^ and RAD51^34^. Therefore, mitochondrial H_2_O_2_ release could affect processes downstream of DNA damage, such as DNA repair.

While superoxide is unlikely to be released from mitochondria due to efficient dismutation, it is a relevant ROS because it can damage iron-sulfur (FeS) clusters, that are assembled in mitochondria^35,36^. Several proteins involved in DNA replication and DNA repair require FeS clusters as cofactors^37,38^, and hence impaired FeS biosynthesis could exacerbate DNA damage stemming from other sources. Iron release could also drive the Fenton reaction mediated ^•^OH production and subsequent damage. Intramitochondrial ROS could also be different than H_2_O_2_ release in the sense that it could potentially damage the mitochondrial genome, which also has been suggested to contribute to cancer development^39^. Using KillerRed to produce superoxide directly in mitochondria instead of H_2_O_2_ could be a complementary approach to DAAO^40^.

In this study we used untransformed RPE1^hTERT^ cells to study the effects of local H_2_O_2_ production. However, genetic alterations like those obtained during cancer progression influence how cells respond to endogenously generated ROS. Genomic changes that lead to an increased reductive capacity and/or a loss of ferroptosis sensitivity^41,42^ are also common adaptations found in cancer cells. For instance, we show that loss of p53 leads to bypass of cell cycle arrest and senescence in response to H_2_O_2_ production at the nucleosome. In addition to genetics, the local metabolomic environment is also strongly linked to the redox properties of a cell^6,14,43^, and can differ greatly between sites in the body or healthy and diseased state. For example, the tumor microenvironment generally provides less nutrients and oxygen, has a lower pH and contains more metabolic waste products^44^, all of which could influence factors like H_2_O_2_ diffusion range and sensitivity to ferroptosis. It is therefore not unthinkable that mitochondria-derived ROS does play a role in the further accumulation of mutations during later stages of tumor progression.

Our findings have possible implications for cancer therapies that aim to boost ROS production to induce cancer cell senescence or death^45^. Our work indicates that ROS-inducing agents might indeed be useful to induce cell cycle arrest and senescence, but that these should be targeted to the nucleus and will likely only be effective in p53-expressing tumors. Ferroptotic cell death induced by cytoplasmic H_2_O_2_, like in RPE1-hTert-DAAO^TOM20^ cells seems to occur largely independent of p53 (Fig S4).

As mentioned, 8-oxo-dG has been detected in patient material, but the source of the oxidant is not known. Based on our observations we envision two scenarios. 1) Guanine in the genomic DNA is subject to oxidation, which would mean that H_2_O_2_ is produced and reacts to form ^•^OH in close proximity to the DNA. A potential source of H_2_O_2_ in the nucleus is for instance Lysin specific demethylase 1 (LSD1), which removes mono- and dimethyl groups on lysine 4 of histone 3 (H3), generating H_2_O_2_ ^46^ and has indeed been shown to lead to the local formation of 8-oxo-dG^47^. NOX4^48^ and the oxidoreductase MICAL1^49^ have also been shown to produce H_2_O_2_ in the nucleus. 2) Free deoxy-GTP nucleotides are oxidized in the cytoplasm followed by incorporation into the genomic DNA. Which of these two scenarios takes place could in principle be distinguished, because free nucleotide incorporation in the DNA gives rise to a different SBS signature compared to direct 8-oxo-dG formation in nuclear DNA^50,51^. SBS signature analysis could also provide a sensitive method to detect sporadic damage and subsequent mutations upon H_2_O_2_ production by localized DAAO, that would stay potentially below a detectable level of DDR activation.

Induction of the ATM/ATR dependent DDR and a subsequent p53-induced cell cycle arrest and senescence observed upon DAAO activity at the nucleosome is reminiscent of the response of RPE1 and other cell types to a bolus of exogenous H_2_O_2_^52,53^. This would suggest that exogenously added H_2_O_2_, unlike H_2_O_2_ produced at the mitochondria in RPE1-hTert-DAAO^TOM20^ cells does make it into the nucleus to induce DNA damage. This apparent discrepancy could potentially be explained by the dose of exogenous H_2_O_2_ that is used in these studies to induce senescence, which was 150-250 µM: much higher than what can be achieved by localized DAAO. Bolus addition of these amounts of H_2_O_2_ lead to overoxidation of PRDX (Fig S1), which may be why it can induce DNA damage. What remains to be elucidated is why these high levels of exogenous H_2_O_2_ do not induce widespread ferroptotic cell death like we observed in response to prolonged exposure of RPE1-hTert-DAAO^TOM20^ cells to >20 mM D-Ala, which does not lead to PRDX overoxidation. Overoxidation of PRDX has been shown to protect yeast cells from cell death upon bolus H_2_O_2_ treatment, and it could be that similar mechanisms underlie our observations^53^.

Taken together, our work shows that not only the dose but also the localization and kinetics of H_2_O_2_ production can have dramatic effects on phenotypic outcome.

## Materials & Methods

### Cell culture

Retinal pigment epithelial cells (hTERT RPE-1, ATCC, CRL-4000) were cultured in DMEM-F12 (Gibco), supplemented with 10% FBS (Bodinco), 100 Units Penicllin-Streptomycin (Merck Life science) and 2 mM L-glutamine (Merck Life science). Cells were grown at 37 °C under 6% CO_2_ atmosphere. Human epithelial kidney cells (HEK293T, ATCC, CRL-3216) used for lentiviral production were cultured in DMEM-High Glucose supplemented with 10% FBS, 100 Units Penicllin-Streptomycin and 2 mM L-glutamine. L-alanine (Merck Life science) and D-alanine (Merck Life science) was dissolved in PBS to create a 1M stock solution and frozen in aliquots, only to be thawed once. Addition of alanine was always precluded by refreshing media. For trapping cells in mitosis to visualize a G1 arrest, 250 ng/ml nocodazole (Merck Life science) was used. For blocking ferroptosis, 500 nM ferrostatin (Merck Life Millipore) or 200 nM Liproxstatin (Merck Life science) was added 1h before treatment with L-or D-Ala.

### Cloning, Lentiviral infection and Cas9 KO procedure

The original genetic sequence for DAAO from *Rhodotorula gracilis* was codon optimized for human cells using the IDT codon optimization tool. The last 3 amino acids containing the localization signal were excluded and the sequence was ordered as a geneblock (Integrated DNA technologies). This construct was a kind gift from Dr. Lucas Bruurs and described before (van Soest et al., manuscript in preparation) The genetic sequence for H2B-mScarlet-I-DAAO and TOM20-mScarlet-I-DAAO was introduced by infusion cloning (Takara Bio) into a lentiviral backbone under control of a human EF-1α promoter and containing a puromycin selection cassette. Plasmids containing the genetic sequence for NLS-HyPer7 and NES-HyPer7 were a kind gift from Dr. Vsevolod Belousov. These sequences were introduced by infusion cloning into a lentiviral backbone under control of a human EF-1α promoter and containing a blasticidin selection cassette. For virus production, lentiviral DAAO and HyPer7 constructs were transiently transfected into HEK293T cells together with 3^rd^ generation packaging vectors using PEI Max (Tebubio). Virus containing media was filtered using a 0.45µm filter and virus was further purified by ultracentrifugation. Virus was added to RPE1-hTERT cells together with 8µg/ml polybrene (Merck Life science). Cells were selected using 5 µg/ml puromycin (Santa Cruz) or 10 µg/ml blasticidin (InvivoGen). Single cells were sorted by FACS and propagated to create monoclonal lines. For the HyPer7 lines, only cells with high HyPer7 expression were sorted.

PCR reactions for cloning were performed using Q5 High-Fidelity DNA polymerase (New England Biolabs), according to manufacturer’s protocol. DNA and plasmid isolation from agarose gels, PCR reactions or bacterial culture was performed using QIAquick Gel extraction kit, QIAprep spin miniprep kit (Qiagen) and Maxiprep purification kit (LabNed).

The construct coding for Cas9, a p53-targeted guide RNA (5’ CGCTATCTGAGCAGCGCTCA 3’) and a GFP expression reporter was made using the PX458 plasmid and method described by *Ran et al*^54^. RPE1-hTert-DAAO^H2B^ and RPE1-hTert-DAAO^TOM20^ were transiently transfected using PEI Max. After 48h GFP positive cells were single-cell sorted into a 96-wells plate using FACS. P53 knock-out status of clones was confirmed by testing sensitivity for the p53 stabilizer Nutlin-3a (10 µM for 16h, Cayman Chemical) and Western Blotting for p53 protein.

### Antibodies

Antibodies used for Western Blot were pChk1 S345 (Cell Signaling, CS2348), Chk1 (Santa Cruz, SC8408), pChk2 T68 (Cell Signaling, CS2661), Chk2 (Cell Signaling, CS3440), γH2AX (Millipore, 05-636), p53 (Santa Cruz, SC-126), p21 (BD Biosciences, 556430), tubulin (Millipore, CP06 OS), prdx2 (Abcam, ab109367) and prdxSO_2/3_ (Abcam, ab16830).

Antibodies used for IF were LaminB1 (Merck Life science, ZRB1141), p21 (Merck Life science, ZRB1141) and γH2AX (Millipore 05-636).

### Live imaging of DAAO localization & Immunofluorescence

Live cell imaging was performed to check whether the mScarlet-I-DAAO fusion proteins localized to the correct subcellular compartments. Cells were grown on glass-bottom dishes (WillcoWells, GWST-3522). Half an hour prior to imaging, media was replaced with media containing 2µg/ml Hoechst33342 (Thermo Fischer Scientific) and 200 nM MitoTracker Green (Thermo Fischer Scientific). After half an hour, media was replaced with fresh media. Cells were imaged on a confocal LSM880 microscope (Zeiss). Hoechst was excited at 405nm and emission measured at 415-481 nm, Mitotracker Green was excited at 488nm and emission measured at 499-545 nm and mScarlet-I was excited at 568nm and measured at 571-677 nm. Z-stacks were acquired and displayed as maximum projections using FIJI software^55^.

For immunofluorescence staining of senescence markers, cells were grown in 8-well Lab-Tek slides with 1.5 borosilicate glass bottom (Thermo Fischer Scientific). Cells were treated for 2 days with L-or D-Alanine and kept in culture for an additional 5 days. Cells were then washed on ice twice with PBS, after which they were fixed with 4% PFA (diluted in PBS from 16% stock, Electron Microscopy Sciences) at room temperature for 15 minutes. Cells were then washed twice with PBS and quenched with 50 mM Glycine (in PBS, Merck Life science) for 10 minutes. Then, cells were twice washed with PBS and permeabilized with 0.1% TX-100 (in PBS, Merck Life science). Cells were washed an additional 2 times, followed by blocking with 2% BSA solution (Merck Life science) in PBS. Primary antibody staining was performed overnight at 4°C in 2% BSA PBS solution containing normal goat IgG (1:10000, Santa-Cruz, 2028). Cells were washed 3x and secondary staining with alexa488 and alexa647 conjugated antibodies (1:500, Life Technologies) was performed at room temp for 1h together with DNA staining by DAPI (1 µg/ml, Merck Life science). Cells were washed an additional 3x and imaged on an LSM880 confocal microscope (Zeiss). Alexa488 was excited at 488nm and emission measured at 493-558nm and Alexa647 excited at 633nm and measured at 638-744nm. Mean signal per nucleus was determined using FIJI software. Nuclei were segmented based on DAPI staining using the StarDist plugin^56^. Mean signal of segmented nuclei was measured.

### Measuring DAAO activity by Oxygen Comsuption Rate

We described a detailed method for the OCR-dependent DAAO activity measurements elsewhere (Manuscript in preparation). Briefly, 24-well V7 Seahorse culture plates were coated with 50µl of 50µg/ml rat tail collagen (Corning, dissolved in 0.1% acetic acid) for 20min at RT, and subsequently washed with PBS. 4*10^4^ DAAO-expressing RPE1^hTERT^ cells were seeded per well. 4 wells remained unseeded for background correction. A prewarmed Seahorse XFe24 analyzer was used to measure OCR. On the day of the assay, the media of the cells was replaced with XF base media supplemented with 2 mM L-Glutamine (Merck Life science), 17.5 mM glucose (Fischer Scientific), 1 mM Na-Pyruvate (Fischer Scientific), and 0.5 mM NaOH. Oligomycin A (final concentration 2µM, Sanbio) and L/D-Alanine was added to injection ports. Oxygen consumption rate was normalized to the third measurement after oligomycin addition.

### HyPer7 measurements

RPE1^hTERT^ cells stably expressing HyPer7 were plated in ibiTreated 8-well µ-slides (Ibidi). Measurements were performed on a Cell Observer microscope (Zeiss) using a 10x magnification. Cells were excited at 385 nm and 475 nm and emission was measured using a 514/44 BP bandpass filter every 10 min for up to 9 hours. After one round of imaging, L/D-Ala was added from 10x concentrated working solutions in PBS to ensure rapid diffusion. Data was quantified using FIJI software^55^. Background of both channels was subtracted in FIJI using the rolling ball algorithm, and channels were thresholded to remove background pixels (NaN). Ratiometric movies of 475_ex_/385_ex_ were made using the image calculator, and mean pixel intensity was measured for each timepoint. Data was normalized first to the L-alanine condition, and then to timepoint zero.

### SDS-PAGE and Western Blot

Protein lysates were obtained by washing cells with PBS and subsequently lysed and scraped in 1x Sample Buffer (2% SDS, 5% 2-mercaptoethanol, 10% glycerol, 0.002% bromophenol blue, 300 mM Tris-HCI pH 6.8,). Samples were first heated for 5 min at 95 °C before running SDS-PAGE and standard blotting procedure. Samples for γH2AX stainings were transferred onto a 0.2 µm nitrocellulose membrane (Cytvia Amersham), whereas other samples were transferred onto a polyvinylidene difluoride (PVDF) membrane (Merck Life science). Samples were blocked for 1h at 4°C in TBS-Tween (1% v/v), containing 1% BSA (Merck Life science). Primary antibody incubation was performed overnight or over weekend at 4°C in TBS-T containing 1% BSA. Secondary antibodies conjugated with horseradish peroxidase (1:10000, Rockland) in TBS-Tween were incubated for 1h at room temperature. Blots were imaged on Image Quant LAS (GE HealthCare).

For non-reducing SDS-PAGE, cells were incubated for 5 min with prewarmed PBS containing 12.5 mg/ml N-Ethylmaleimide (Merck Life science) at 37 °C prior to harvesting to fix the cysteine-dependent redox state. Cells were then washed with PBS and lysed in buffer containing 50 mM Tris-HCl (pH 7.5), 0.1% TritonX-100, 2.25 mM MgCl_2_, 0.1M NaCl, NaF, 0.1% Aprotinin, 0.1% Leupeptin and 100 mM iodoacetamide (all Merck Life science). Samples were divided over two tubes and a reducing and a non-reducing sample was prepared by adding SDS sample buffer with or without beta-mercaproethanol.

### Flow cytometry

Cell viability was assessed by propidium iodide (PI) exclusion. Media and subsequent wash with PBS was collected, after which cells were trypsinized, resuspended and added to the collected medium+PBS (15ml falcon tubes) and spun down (1500 RPM 5min, Beckmann tabletop centrifuge with swingout rotor) and put on ice. Supernatant was removed, and the pelleted cells were resuspended in PBS containing 20µg/ml propium iodide (Merck Life science).

For DNA profiles, cells were collected in a similar fashion. After centrifugation and removal of supernatant, 200 µl of PBS was added to prevent cells from aggregating during fixation. Cells were fixed by dropwise addition of 5ml of ice-cold 70% ethanol while vortexing. Samples were left at 4°C overnight or longer. On the day of measurement, fixed cells were pelleted (1500 RPM 5 min) and resuspended in PBS containing 20 µg/ml propium iodide and 200 µg/ml RNase (Merck Life science).

For BrdU measurements, cells were cultured in media containing 10 µM BrdU (BD Biosciences) 24h prior to collecting cells. Cells were collected and fixed in the same fashion as for DNA profiles. On the day of measurement, anti-BrdU staining was performed. First, after centrifugation the cell pellet was resuspended in 0.1 M HCL solution containing 0.5 mg/ml of the protease pepsin (Merck Life science) and left for 20 min at room temperature. After washing with PBS containing 0.1% BSA and 0.5% Tween, cells were pelleted and treated with 2 M HCL for 12 min at 37°C to denature the DNA, after which double the volume of borate buffer (0.1M H_3_BO_3_ set at pH 8.5 with 0.1M Na_2_B_4_O_7_) was added. After washing and centrifugation, the cell pellet was incubated with PBS/BSA/Tween containing BrdU-FITC antibody (1:50, BD Biosciences) for 1h on ice in the dark. Cells were centrifuged and resuspended in PBS/BSA containing 20 µg/ml PI and 200 µg/ml RNase.

Flow cytometry samples were measured on BD FACSCelesta Cell analyzer (BD biosciences) and analyzed with BD FacsDiva Software.

### Crystal violet staining

Cells were plated in 6-well plates. After 1 day, cells were treated with L-or D-Alanine for 24 hours. Cells were then washed 2x with PBS to remove dead cells, and fixed with ice-cold (-20 °C) methanol for 5 min. Methanol was removed and replaced with crystal violet solution (0.5% w/v in 25% v/v methanol in H_2_O) for 10 min. Crystal violet was removed and wells are washed with water and airdried overnight. Plates were imaged on an Epson 3170 photo scanner (Epson).

### Senescence-associated β-galactosidase staining

For the SA-β-gal staining, cells were cultured in 6-well plates. Cells were treated for 2 days with L-or D-Ala and kept in culture for an additional 5 days. Then cells were washed twice with PBS and fixed for 15 min with 3.7% formalin (37% formalin diluted in PBS). Cells were washed twice with PBS and then incubated with freshly made SA-β-gal buffer containing 25 mM Na_2_HPO_4_ (Merck Life science), 7.4 mM Citric Acid (Merck Life science), 150 mM NaCl (Merck Life science), 2 mM MgCl_2_ (Merck Life science), 5 mM Potassium Ferricyanide (Merck Life science), 5 mM Potassium Ferrocyanide (Merck Life science) and 0.1% X-Gal (from a 4% X-Gal solution in dimethylformamide, Merck Life science). Buffer was passed through a 0.2 micron syringe filter before X-Gal was added. Cells were incubated overnight at 37 °C, at ambient CO_2_. Cells were rinsed 2x with PBS and kept under 70% ethanol. Cells were imaged on an EVOS-M5000 microscope (Thermo Fisher).

### Comet assay

Alkaline comet assays were performed using the CometAssay single cell gel electrophoresis assay kit (R&D systems) according to the manufacturers’ instructions. Cells were treated for 2 hours with L/D-Ala after which they were trypsinized and counted to make 10^5^ cells/ml dilutions. Cell lysis was performed overnight, DNA alkaline unwinding for 1h and electrophoresis for 1h, all at 4°C. DNA staining was performed using Midori Green Advance for 30 min at room temperature (1:5000 in TE buffer pH7.5, Nippon Genetics). Comets were imaged on an EVOS-M5000 microscope (Thermo Fisher) using the GFP filter block. Comets were manually analyzed using FIJI software. Samples were blinded before quantification. The area and mean pixel value of the comet head and the complete comet were measured. In addition, the mean pixel value of the local background was determined. The mean pixel value of the background was subtracted of the mean pixel value of the comet head and total comet. These values were then multiplied with the area of the comet head or total comet to create the integrated densities corrected for background signal. The integrated density of the comet tail value was calculated by subtracting the integrated density of the comet head from the integrated density of the total comet. The percentage DNA in comet tail was then calculated by dividing the integrated density of the comet tail value with the integrated density of the total comet. Per condition 50 comets were analyzed.

## Acknowledgements

We would like to thank Tessa Vreeman for help with constructing lentiviral HyPer7 plasmids. We are grateful to dr. Peter de Keizer and dr. Johannes Lehmann for advise on the assessment of senescence, Livio Kleij for technical support with the HyPer7 measurements, and our colleagues at the Center for Molecular Medicine, UMC Utrecht for their valuable input and suggestions. This work was funded by a grant from the Dutch Cancer Society (KWF Kankerbestrijding) to TBD. BMTB is a member of the Oncode Institute, which is partly funded by the Dutch Cancer Society (KWF Kankerbestrijding).

## References

1 Hanahan, D. & Weinberg, R. A. Hallmarks of cancer: the next generation. Cell 144, 646–674, doi:10.1016/j.cell.2011.02.013 (2011).

2 Lopez-Otin, C., Blasco, M. A., Partridge, L., Serrano, M. & Kroemer, G. Hallmarks of aging: An expanding universe. Cell 186, 243–278, doi:10.1016/j.cell.2022.11.001 (2023).

3 Krokan, H. E. & Bjoras, M. Base excision repair. Cold Spring Harb Perspect Biol 5, a012583, doi:10.1101/cshperspect.a012583 (2013).

4 Kurthkoti, K., Kumar, P., Sang, P. B. & Varshney, U. Base excision repair pathways of bacteria: new promise for an old problem. Future Med Chem 12, 339–355, doi:10.4155/fmc-2019-0267 (2020).

5 Grasso, S. & Tell, G. Base excision repair in Archaea: back to the future in DNA repair. DNA Repair (Amst) 21, 148–157, doi:10.1016/j.dnarep.2014.05.006 (2014).

6 Sies, H. & Jones, D. P. Reactive oxygen species (ROS) as pleiotropic physiological signalling agents. Nat Rev Mol Cell Biol 21, 363–383, doi:10.1038/s41580-020-0230-3 (2020).

7 Cadet, J., Douki, T. & Ravanat, J. L. Oxidatively generated base damage to cellular DNA. Free Radic Biol Med 49, 9–21, doi:10.1016/j.freeradbiomed.2010.03.025 (2010).

8 Aruoma, O. I., Halliwell, B. & Dizdaroglu, M. Iron ion-dependent modification of bases in DNA by the superoxide radical-generating system hypoxanthine/xanthine oxidase. J Biol Chem 264, 13024–13028 (1989).

9 Aruoma, O. I., Halliwell, B., Gajewski, E. & Dizdaroglu, M. Damage to the bases in DNA induced by hydrogen peroxide and ferric ion chelates. J Biol Chem 264, 20509–20512 (1989).

10 Murphy, M. P. How mitochondria produce reactive oxygen species. Biochem J 417, 1–13, doi:10.1042/BJ20081386 (2009).

11 Barnes, D. E. & Lindahl, T. Repair and genetic consequences of endogenous DNA base damage in mammalian cells. Annu Rev Genet 38, 445–476, doi:10.1146/annurev.genet.38.072902.092448 (2004).

12 White, E. Deconvoluting the context-dependent role for autophagy in cancer. Nat Rev Cancer 12, 401–410, doi:10.1038/nrc3262 (2012).

13 Yang, Y. et al. Mitochondria and Mitochondrial ROS in Cancer: Novel Targets for Anticancer Therapy. J Cell Physiol 231, 2570–2581, doi:10.1002/jcp.25349 (2016).

14 Hoehne, M. N. et al. Spatial and temporal control of mitochondrial H2 O2 release in intact human cells. EMBO J 41, e109169, doi:10.15252/embj.2021109169 (2022).

15 Pak, V. V. et al. Ultrasensitive Genetically Encoded Indicator for Hydrogen Peroxide Identifies Roles for the Oxidant in Cell Migration and Mitochondrial Function. Cell Metab 31, 642–653 e646, doi:10.1016/j.cmet.2020.02.003 (2020).

16 Cleaver, J. E. et al. Mitochondrial reactive oxygen species are scavenged by Cockayne syndrome B protein in human fibroblasts without nuclear DNA damage. Proc Natl Acad Sci U S A 111, 13487–13492, doi:10.1073/pnas.1414135111 (2014).

17 Shi, T., van Soest, D. M. K., Polderman, P. E., Burgering, B. M. T. & Dansen, T. B. DNA damage and oxidant stress activate p53 through differential upstream signaling pathways. Free Radic Biol Med 172, 298–311, doi:10.1016/j.freeradbiomed.2021.06.013 (2021).

18 Kirova, D. G. et al. A ROS-dependent mechanism promotes CDK2 phosphorylation to drive progression through S phase. Dev Cell 57, 1712–1727 e1719, doi:10.1016/j.devcel.2022.06.008 (2022).

19 Steinhorn, B. et al. Chemogenetic generation of hydrogen peroxide in the heart induces severe cardiac dysfunction. Nat Commun 9, 4044, doi:10.1038/s41467-018-06533-2 (2018).

20 Erdogan, Y. C. et al. Complexities of the chemogenetic toolkit: Differential mDAAO activation by d-amino substrates and subcellular targeting. Free Radic Biol Med 177, 132–142, doi:10.1016/j.freeradbiomed.2021.10.023 (2021).

21 Brand, M. D. The sites and topology of mitochondrial superoxide production. Exp Gerontol 45, 466–472, doi:10.1016/j.exger.2010.01.003 (2010).

22 Moller, M. N. et al. Diffusion and Transport of Reactive Species Across Cell Membranes. Adv Exp Med Biol 1127, 3–19, doi:10.1007/978-3-030-11488-6_1 (2019).

23 Wang, Y., Branicky, R., Noe, A. & Hekimi, S. Superoxide dismutases: Dual roles in controlling ROS damage and regulating ROS signaling. J Cell Biol 217, 1915–1928, doi:10.1083/jcb.201708007 (2018).

24 Gorini, F., Scala, G., Cooke, M. S., Majello, B. & Amente, S. Towards a comprehensive view of 8-oxo-7,8-dihydro-2’-deoxyguanosine: Highlighting the intertwined roles of DNA damage and epigenetics in genomic instability. DNA Repair (Amst) 97, 103027, doi:10.1016/j.dnarep.2020.103027 (2021).

25 Stallaert, W. et al. The molecular architecture of cell cycle arrest. Mol Syst Biol 18, e11087, doi:10.15252/msb.202211087 (2022).

26 Feringa, F. M. et al. Persistent repair intermediates induce senescence. Nat Commun 9, 3923, doi:10.1038/s41467-018-06308-9 (2018).

27 Feringa, F. M. et al. Hypersensitivity to DNA damage in antephase as a safeguard for genome stability. Nat Commun 7, 12618, doi:10.1038/ncomms12618 (2016).

28 Hornsveld, M. et al. A FOXO-dependent replication checkpoint restricts proliferation of damaged cells. Cell Rep 34, 108675, doi:10.1016/j.celrep.2020.108675 (2021).

29 Narita, M. et al. Rb-mediated heterochromatin formation and silencing of E2F target genes during cellular senescence. Cell 113, 703–716, doi:10.1016/s0092-8674(03)00401-x (2003).

30 Alexandrov, L. B. et al. Signatures of mutational processes in human cancer. Nature 500, 415–421, doi:10.1038/nature12477 (2013).

31 Morgan, B. et al. Real-time monitoring of basal H2O2 levels with peroxiredoxin-based probes. Nat Chem Biol 12, 437–443, doi:10.1038/nchembio.2067 (2016).

32 Kang, D. et al. Intracellular localization of 8-oxo-dGTPase in human cells, with special reference to the role of the enzyme in mitochondria. J Biol Chem 270, 14659–14665, doi:10.1074/jbc.270.24.14659 (1995).

33 Guo, Z., Kozlov, S., Lavin, M. F., Person, M. D. & Paull, T. T. ATM activation by oxidative stress. Science 330, 517–521, doi:10.1126/science.1192912 (2010).

34 Skoko, J. J. et al. Redox regulation of RAD51 Cys319 and homologous recombination by peroxiredoxin 1. Redox Biol 56, 102443, doi:10.1016/j.redox.2022.102443 (2022).

35 Lill, R. & Freibert, S. A. Mechanisms of Mitochondrial Iron-Sulfur Protein Biogenesis. Annu Rev Biochem 89, 471–499, doi:10.1146/annurev-biochem-013118-111540 (2020).

36 Jang, S. & Imlay, J. A. Micromolar intracellular hydrogen peroxide disrupts metabolism by damaging iron-sulfur enzymes. J Biol Chem 282, 929–937, doi:10.1074/jbc.M607646200 (2007).

37 Netz, D. J. et al. Eukaryotic DNA polymerases require an iron-sulfur cluster for the formation of active complexes. Nat Chem Biol 8, 125–132, doi:10.1038/nchembio.721 (2011).

38 Fuss, J. O., Tsai, C. L., Ishida, J. P. & Tainer, J. A. Emerging critical roles of Fe-S clusters in DNA replication and repair. Biochim Biophys Acta 1853, 1253–1271, doi:10.1016/j.bbamcr.2015.01.018 (2015).

39 Smith, A. L. M. et al. Author Correction: Age-associated mitochondrial DNA mutations cause metabolic remodeling that contributes to accelerated intestinal tumorigenesis. Nat Cancer 2, 129, doi:10.1038/s43018-020-00156-7 (2021).

40 Koren, S. A. et al. All-optical spatiotemporal mapping of ROS dynamics across mitochondrial microdomains <em>in situ</em>. bioRxiv, 2023.2001.2007.523093, doi:10.1101/2023.01.07.523093 (2023).

41 Cheung, E. C. & Vousden, K. H. The role of ROS in tumour development and progression. Nat Rev Cancer 22, 280–297, doi:10.1038/s41568-021-00435-0 (2022).

42 Hornsveld, M. & Dansen, T. B. The Hallmarks of Cancer from a Redox Perspective. Antioxidants & redox signaling 25, 300–325 (2016).

43 Panieri, E. & Santoro, M. M. ROS homeostasis and metabolism: a dangerous liason in cancer cells. Cell Death Dis 7, e2253, doi:10.1038/cddis.2016.105 (2016).

44 Lyssiotis, C. A. & Kimmelman, A. C. Metabolic Interactions in the Tumor Microenvironment. Trends Cell Biol 27, 863–875, doi:10.1016/j.tcb.2017.06.003 (2017).

45 Wondrak, G. T. Redox-directed cancer therapeutics: molecular mechanisms and opportunities. Antioxid Redox Signal 11, 3013–3069, doi:10.1089/ars.2009.2541 (2009).

46 Perillo, B., Tramontano, A., Pezone, A. & Migliaccio, A. LSD1: more than demethylation of histone lysine residues. Exp Mol Med 52, 1936–1947, doi:10.1038/s12276-020-00542-2 (2020).

47 Perillo, B. et al. DNA oxidation as triggered by H3K9me2 demethylation drives estrogen-induced gene expression. Science 319, 202–206, doi:10.1126/science.1147674 (2008).

48 Martyn, K. D., Frederick, L. M., von Loehneysen, K., Dinauer, M. C. & Knaus, U. G. Functional analysis of Nox4 reveals unique characteristics compared to other NADPH oxidases. Cell Signal 18, 69–82, doi:10.1016/j.cellsig.2005.03.023 (2006).

49 Fremont, S. et al. Oxidation of F-actin controls the terminal steps of cytokinesis. Nat Commun 8, 14528, doi:10.1038/ncomms14528 (2017).

50 Suzuki, T. & Kamiya, H. Mutations induced by 8-hydroxyguanine (8-oxo-7,8-dihydroguanine), a representative oxidized base, in mammalian cells. Genes Environ 39, 2, doi:10.1186/s41021-016-0051-y (2017).

51 Satou, K., Kawai, K., Kasai, H., Harashima, H. & Kamiya, H. Mutagenic effects of 8-hydroxy-dGTP in live mammalian cells. Free Radic Biol Med 42, 1552–1560, doi:10.1016/j.freeradbiomed.2007.02.024 (2007).

52 Putker, M. et al. Redox-dependent control of FOXO/DAF-16 by transportin-1. Mol Cell 49, 730–742, doi:10.1016/j.molcel.2012.12.014 (2013).

53 Marazita, M. C., Dugour, A., Marquioni-Ramella, M. D., Figueroa, J. M. & Suburo, A. M. Oxidative stress-induced premature senescence dysregulates VEGF and CFH expression in retinal pigment epithelial cells: Implications for Age-related Macular Degeneration. Redox Biol 7, 78–87, doi:10.1016/j.redox.2015.11.011 (2016).

54 Ran, F. A. et al. Genome engineering using the CRISPR-Cas9 system. Nat Protoc 8, 2281–2308, doi:10.1038/nprot.2013.143 (2013).

55 Schindelin, J. et al. Fiji: an open-source platform for biological-image analysis. Nat Methods 9, 676–682, doi:10.1038/nmeth.2019 (2012).

56 Schmidt, U., Weigert, M., Broaddus, C. & Myers, G. in Medical Image Computing and Computer Assisted Intervention – MICCAI 2018. (eds Alejandro F. Frangi et al.) 265–273 (Springer International Publishing).

