## Supplemental figures for "Mitochondrial H_2_O_2_ release does not directly cause genomic DNA damage"

A.

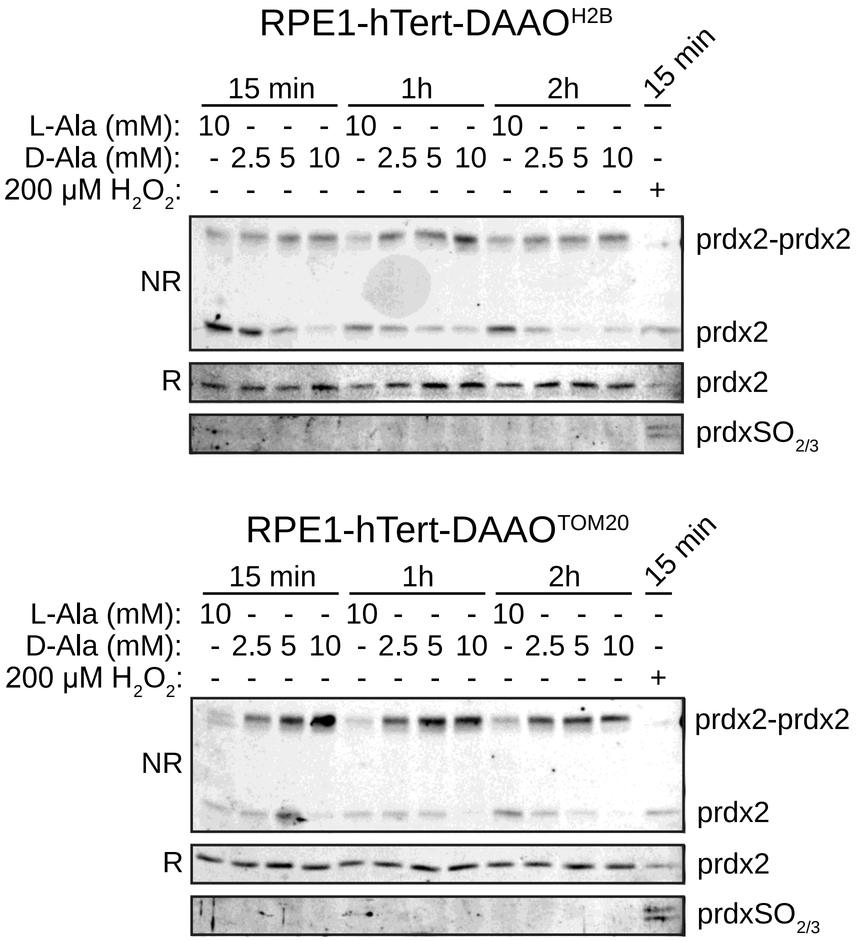


B.

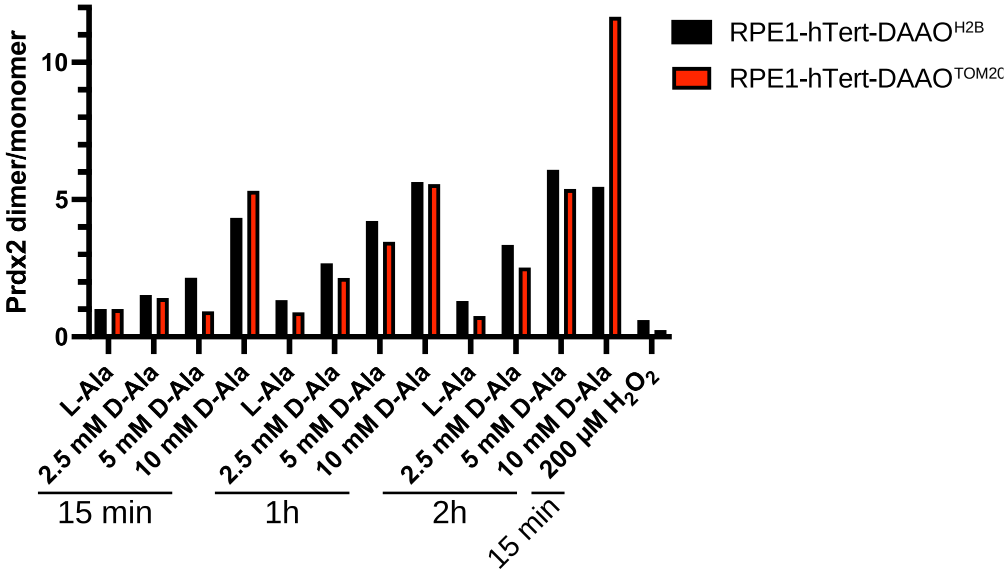


Sup.Fig 1: DAAO-produced $H_2O_2$ induces prdx2 dimerization but no overoxidation

(A) Non-reducing (NR) & Reducing (R) western blots for prdx2 and prdxSO$_{2/3}$ in RPE1-hTert-DAAO$^{H2B}$ and RPE1-hTert-DAAO$^{TOM20}$ cells. DAAO activation by D-Alanine results in prdx2

oxidation and subsequent dimerization in a concentration dependent manner, but does not result in prdx overoxidation (prdxSO$_{2/3}$), in contrast to exogenous $H_2O_2$ treatment.
(B) Quantification of the non-reducing prdx2 blot shown in A, showing the prdx2 dimer over monomer intensity ratio. Values are normalized to 15 min L-Ala condition.

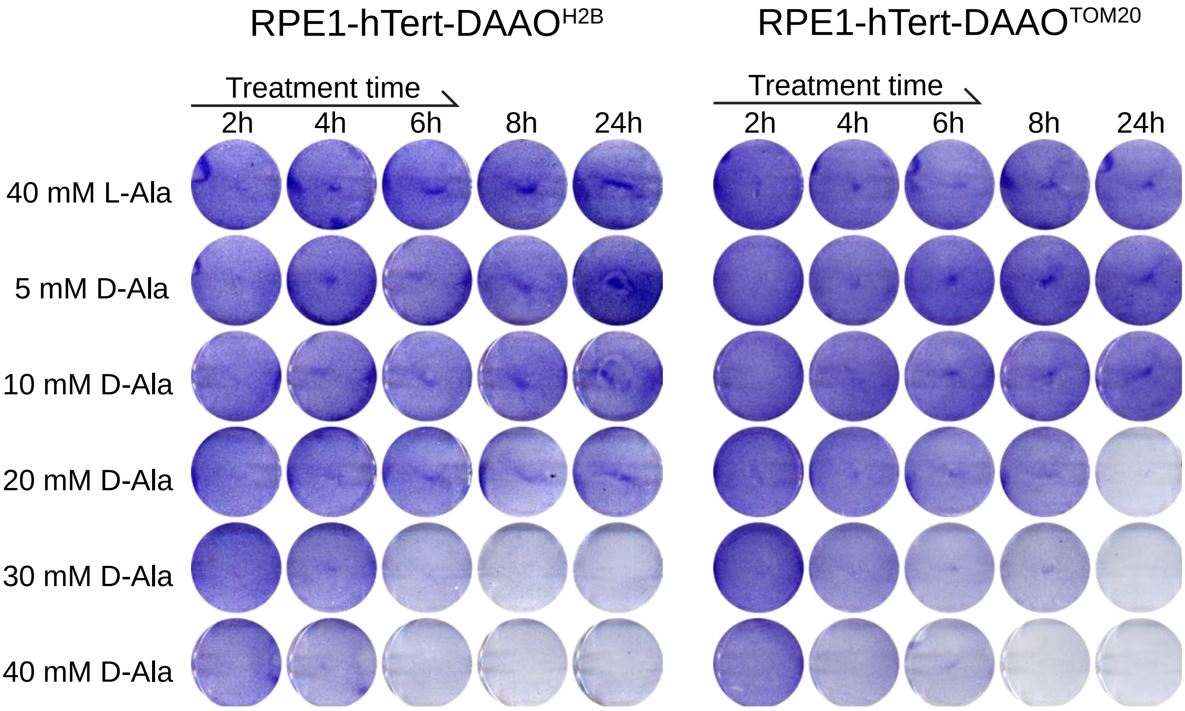


<u>Sup. fig. 2: Viability is affected by both DAAO activity as well as duration of DAAO activation</u>

Crystal Violet staining of RPE1-hTert-DAAO$^{H2B}$ and RPE1-hTert-DAAO$^{TOM20}$ treated with several concentrations of L/D-Ala for different times. Cells were fixed and stained 24h after treatment was initiated. Viability does not only decrease by increasing DAAO activity but also by extending the treatment duration.

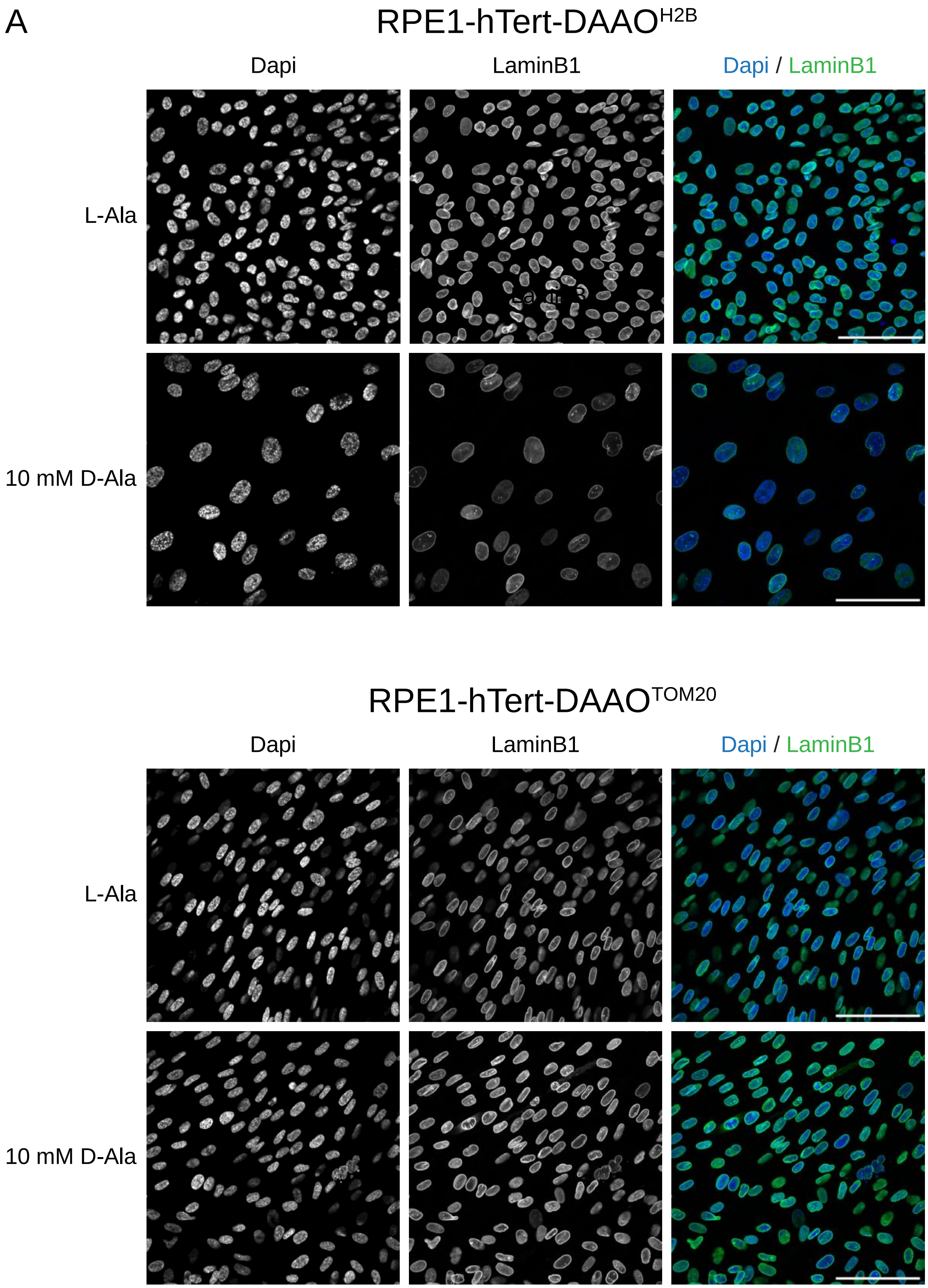
A
RPE1-hTert-DAAO$^{H2B}$
Dapi
LaminB1
Dapi / LaminB1
L-Ala
10 mM D-Ala
RPE1-hTert-DAAO$^{TOM20}$
Dapi
LaminB1
Dapi / LaminB1
L-Ala
10 mM D-Ala

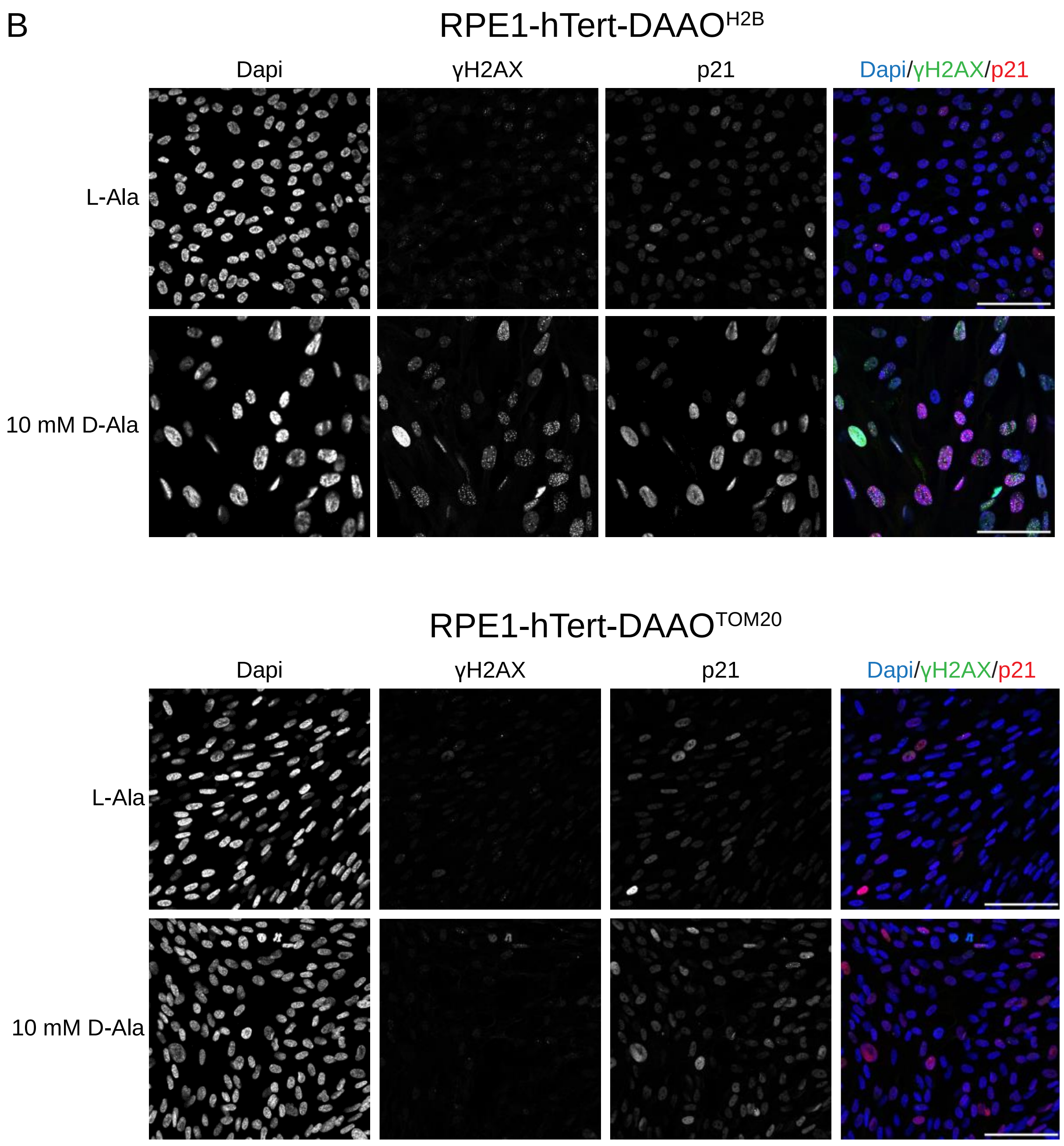


Sup. Figure 3: RPE1 cells display senescence markers upon $H_2O_2$ production at nucleosomes

IF pictures of senescence markers in RPE1-hTert-DAAO$^{H2B}$ and RPE1-hTert-DAAO$^{TOM20}$ cells (scale bar = 100 µm). Cells develop decreased laminB1 (A) levels and increased p21 expression (B) upon $H_2O_2$ production at the nucleosomes. They also display increased γH2AX foci, indicative of DNA strand breaks. This is not observed when mitochondrial $H_2O_2$ release is mimicked in RPE1-hTert-DAAO$^{TOM20}$ cells.

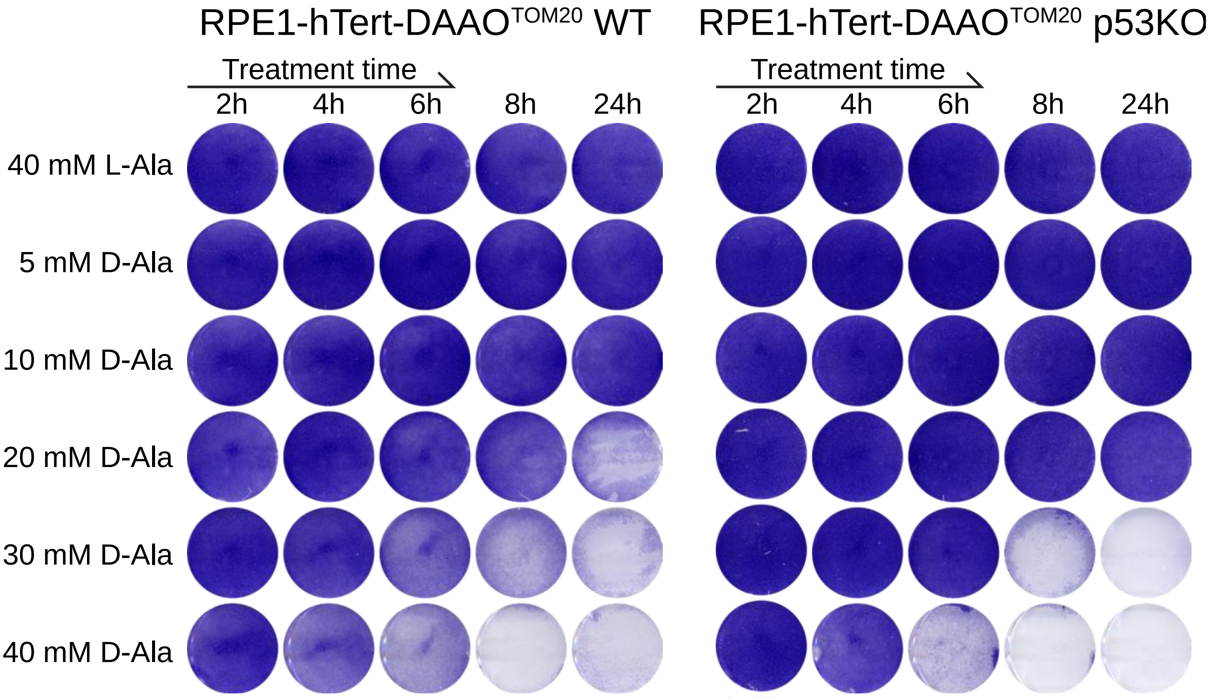


Sup. fig. 4: Ferroptosis induction in RPE1-hTert-DAAO$^{TOM20}$ cells is not p53 dependent.
Crystal Violet staining of RPE1-hTert-DAAO$^{TOM20}$ WT and p53KO cells treated with several concentrations of L/D-Ala for different times. Cells were fixed and stained 24h after treatment was initiated. Induction of ferroptosis seems to be largely independent of p53 status.
